# Single-cell chromatin state transitions during epigenetic memory formation

**DOI:** 10.1101/2023.10.03.560616

**Authors:** Taihei Fujimori, Carolina Rios-Martinez, Abby R. Thurm, Michaela M. Hinks, Benjamin R. Doughty, Joydeb Sinha, Derek Le, Antonina Hafner, William J. Greenleaf, Alistair N. Boettiger, Lacramioara Bintu

**Affiliations:** Department of Bioengineering, Stanford University, Stanford, CA, USA; Biophysics Program, Stanford University, Stanford, CA, USA; Department of Genetics, Stanford University, Stanford, CA, USA; Department of Chemical & Systems Biology, Stanford University, Stanford, CA, USA; Department of Dermatology, Program in Epithelial Biology, Stanford University, Stanford, CA, USA; Program in Cancer Biology, Stanford University, Stanford, CA, USA; Department of Developmental Biology, Stanford University, Stanford, CA, USA; Department of Discovery Oncology, Genentech, CA, USA; Department of Applied Physics, Stanford University, Stanford, CA, USA; Chan Zuckerberg Biohub, San Francisco, CA, USA

## Abstract

Repressive chromatin modifications are thought to compact chromatin to silence transcription. However, it is unclear how chromatin structure changes during silencing and epigenetic memory formation. We measured gene expression and chromatin structure in single cells after recruitment and release of repressors at a reporter gene. Chromatin structure is heterogeneous, with open and compact conformations present in both active and silent states. Recruitment of repressors associated with epigenetic memory produces chromatin compaction across 10-20 kilobases, while reversible silencing does not cause compaction at this scale. Chromatin compaction is inherited, but changes molecularly over time from histone methylation (H3K9me3) to DNA methylation. The level of compaction at the end of silencing quantitatively predicts epigenetic memory weeks later. Similarly, chromatin compaction at the Nanog locus predicts the degree of stem-cell fate commitment. These findings suggest that the chromatin state across tens of kilobases, beyond the gene itself, is important for epigenetic memory formation.

## Introduction

Chromatin modifications on both DNA and histone tails are a fundamental aspect of eukaryotic gene regulation. It is widely thought that repressive chromatin modifications are associated with chromatin compaction^1^. Indeed, histone H3 lysine 9 trimethylation (H3K9me3), one of the main repressive modifications, is associated with compacted heterochromatin regions^2^. Targeted recruitment of the KRAB domain from Zinc Finger 10 to a gene causes silencing^3,4^ through its direct interaction with KAP1, a corepressor that in turn recruits histone methylase SETDB1 (the writer of H3K9me3), histone deacetylases, and heterochromatin protein 1 (HP1, the reader of H3K9me3)^5,6^. Both KRAB and HP1 recruitment at a genomic locus induce de novo accumulation of H3K9me3, increase chromatin density^7^ and reduce chromatin accessibility^8^. Recruitment of dCas9-KRAB (known as CRISPRi^4^) to many loci reorganizes chromatin into a heterochromatin-like structure^9^.

These heterochromatin-associated modifications are often transmitted across multiple cell divisions through positive feedback mediated by reader-writer complexes, and are connected with epigenetic memory^10,11^. Epigenetic memory is important for converting transient perturbations, such as differentiation signals^12^, infections^13^, or metabolic stresses^14^ into heritable gene expression response programs. By using a synthetic approach, it has been shown that targeted KRAB recruitment and release from a reporter gene causes a fraction of cells to remain stably silenced, while the rest reactivate gene expression^15^. The fraction of individual cells that display this all-or-none epigenetic memory can be modulated by varying the duration of recruitment^15^; thus this synthetic recruitment system constitutes an ideal model for tuning epigenetic memory. Moreover, this fractional response suggests heterogeneity of chromatin states within single cells, which could be associated with heterogeneous 3D chromatin structures, similarly to recent observations from chromatin imaging in single cells in other contexts^16,17^.

While it is clear that repressive chromatin modifications and compacted heterochromatin are correlated with silent genes, it is still not clear how dynamic changes in chromatin structure and molecular state connect to gene silencing and memory formation: How heterogeneous is chromatin structure at the single-cell level? How does chromatin structure and chromatin modifications change at a repressor-targeted locus and its neighborhood (tens of kilobases surrounding it)? Does chromatin state change as cells commit from silencing to epigenetic memory and does it affect memory formation? To answer these questions, we combined the targeted recruitment and release of repressors that allows us to tune silencing and epigenetic memory^15^ with a multiplexed DNA FISH method for measuring chromatin structure in single cells^18^. We found that KRAB recruitment, known to cause epigenetic memory, leads to H3K9me3 accumulation and chromatin compaction across tens of kilobases. Compaction arises at the population average level, but chromatin structure is heterogeneous in single cells, with open and compacted conformations present in both active and silent cells. The compaction is retained in stably silenced cells even after KRAB release, despite the fact that H3K9me3 histone methylation is lost and replaced by DNA methylation. In contrast, silencing by histone deacetylase HDAC4 does not lead to epigenetic memory nor large-scale compaction, suggesting transcriptional silencing is not sufficient to induce chromatin compaction at this length scale. By varying the duration of KRAB recruitment and using KRAB mutants with partial loss of function, we generated cell populations with different percentages of stably silenced cells, and found that the degree of chromatin compaction across 10-20 kb at the end of recruitment can quantitatively predict epigenetic memory weeks later. This chromatin compaction at the end of recruitment is associated with spreading of H3K9me3, especially across regions enriched in H3K9 acetylation before silencing. Finally, to determine whether the connection between compaction and epigenetic memory affects important biological processes such as cell fate commitment, we measured compaction of the Nanog locus, a hallmark of pluripotency, as it undergoes silencing during mouse stem cell differentiation. We found that compaction at the Nanog locus is a good predictor of fate commitment quantified by the number of cells that cannot revert to the pluripotent state after withdrawal of the differentiation stimulus, suggesting that chromatin compaction upon epigenetic silencing is predictive of irreversible fate commitment.

## Results

### Single-cell chromatin tracing after chromatin regulator recruitment and release

We developed a system where we can control gene silencing and epigenetic memory by recruitment of a chromatin regulator and measure changes in chromatin structure at the target locus and the region surrounding it. We used a synthetic reporter system (Fig. 1a) consisting of a fluorescent protein, mCitrine, expressed under the control of a constitutive human EF1alpha promoter with 9x TetO recruitment sites^19^. The reporter construct was stably integrated into the AAVS1 safe harbor site on chromosome 19 in immortalized human embryonic kidney cells (HEK293T). The KRAB repressor domain from ZNF10 was fused to the reverse tetracycline repressor (rTetR)^20^, facilitating precise control of their recruitment and release through addition and removal of doxycycline (dox) to the culture media. After 5 days of KRAB recruitment, which results in H3K9me3 accumulation (Extended Data Fig. 1a), as reported before ^7,21^, we observe complete silencing of the reporter (Fig. 1b, middle). Releasing KRAB from the target gene results in reactivation of gene expression in a subset of cells while the rest remain silenced, leading to a bimodal distribution of reporter expression levels which reaches steady state within 12 days after release (Fig. 1b right, Fig. 1c), consistent with the all-or-none epigenetic memory formation previously described^15^. Moreover, after dox removal and cell sorting, the silenced and reactivated states were stable over time, indicating that a fraction of cells become irreversibly silenced after KRAB recruitment (Fig. 1d).

**Fig. 1.**
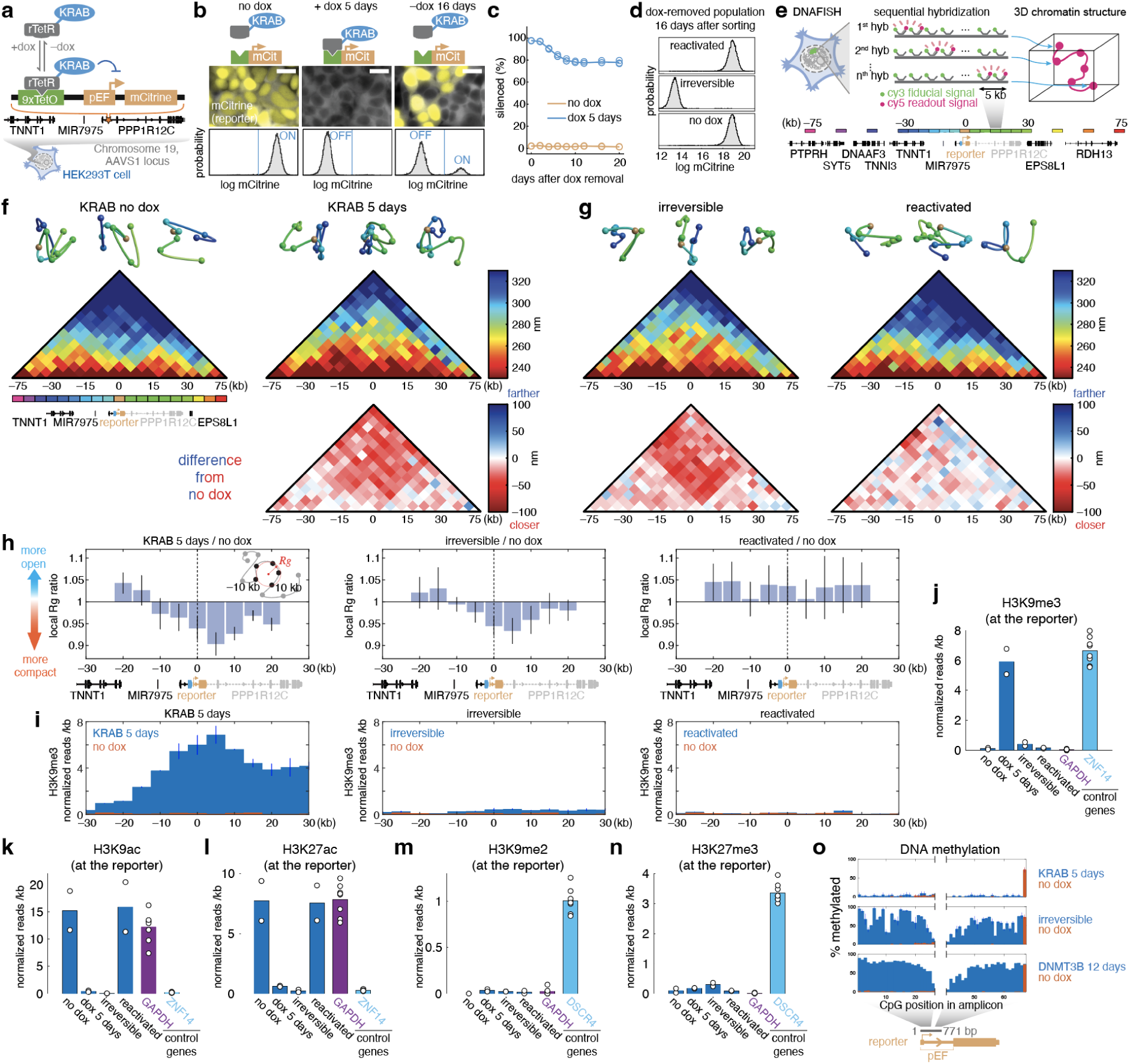
Targeted chromatin regulator recruitment results in 3D chromatin compaction that is retained in irreversibly silenced cells. (a) Schematic representation of the reporter system with targeted chromatin regulator recruitment. Reporter (pEF-mCitrine) orange with 9 TetO sites (green), allowing recruitment of rTetR-KRAB (blue) by dox addition, is integrated HEK293T cells at the AAVS1 locus in the PPP1R12C gene (top). (b) Reporter gene expression in response to KRAB recruitment (A6 clone). (top) Fluorescent images show mCherry co-expressed with rTetR-KRAB (gray) and the reporter mCitrine (yellow) in no dox (left), +dox for 5 days (center), dox removed for 16 days after the 5 day recruitment (right). Scale bar: 20 µm. (bottom) Probability density distributions for mCitrine fluorescence measured by flow cytometry. (c) Time-course showing the percentage of cells with mCitrine silenced (OFF population in b) for the A6 cell lines during the KRAB release period following 5 days of KRAB recruitment (blue) or no dox control (yellow). Lines show the average of three replicates, circles show each replicate. (d) Distributions of mCitrine expression levels in the irreversibly silenced or reactivated cells following cell sorting (B9 clone). (e) (top) Schematic overview of the ORCA technique. (bottom) Genome segments that are imaged (5kb each) are annotated with colored rectangles above the genome browser view of the locus. (f) (top) Representative 3D chromatin structure (from -30 kb to 30 kb) without dox treatment (topleft) or with 1 µg/mL dox treatment for 5 days to recruit rTetR-KRAB (topright). Color-code follows the schematic in Fig. 1d, with the reporter region in orange. Detected segments (spheres) are connected with spline curves for visualization. (middle) Heatmaps of median distances between pairwise detected segments. (bottom) Subtracted median distance maps upon KRAB recruitment compared to the no-dox control, where red colors indicate shorter distances. Data from five replicates (Extended Data Fig. 6a,b) are averaged. (g) (top) Representative 3D chromatin structure, (middle) median distance maps and (bottom) subtracted median distance maps in (left) irreversibly silenced or (right) reactivated cells after 1 µg/mL dox treatment for 5 days and subsequent dox washout (for >16 days). Data from four or six replicates (Extended Data Fig. 6a,b) are averaged. (h) (left) The local radius of gyration (Rg) is quantified at each genomic position using 5 consecutive segments (±10 kb). (right) The ratio of the local radius of gyration compared to no dox control from the same sample. Mean ± s.e.m. (i) 5 kb-binned H3K9me3 CUT&RUN profile (B9 clone). Data from two replicates are averaged. Vertical bars show maximum and minimum values. (j-n) Quantification of normalized CUT&RUN reads at the reporter integrated region with antibodies against: (j) H3K9me3, (k) H3K9ac, (l) H3K27ac, (m) H3K9me2, (n) H3K27me3 from cells in the indicated conditions (no dox, dox for 5 days, irreversibly silenced, and reactivated). Reads at the pEF promoter region are excluded from the H3K9ac and H3K27ac quantifications, since for these modifications reads also align at the endogenous pEF1 alpha copies in addition to the reporter one (Methods). All CUT&RUN experiments were done with the B9 clone (see Extended Data Fig. 4). Bars show the average of two replicates, circles show each replicate. Normalized reads of positive and negative control genes averaged across all conditions and replicates are shown with dots that represent each replicate. (o) Quantification of DNA methylation (% methylated CpG sites) at the reporter measured by EM-seq in the following conditions: (top) after KRAB recruitment for 5 days, (middle) in cells irreversibly silenced by KRAB, (bottom) after DNMT3B recruitment for 12 days as a positive control for DNA methylation. Bars show data averaged from two replicates of each experimental condition (blue) or no dox control (orange). Lines show minimum and maximum values.

To visualize chromatin structure in single cells, we used a microscopy chromatin tracing technique called ORCA (Optical Reconstruction of Chromatin Architecture^18^). ORCA relies on sequential hybridization and imaging steps in order to measure distances in units of nm between all points along the chromatin trace (Fig. 1e). In order to perform automated rounds of fluorescent probe hybridization, we added a custom-built liquid handling system to a commercial wide-field epifluorescence microscope (Methods). To validate the performance of this system relative to the original ORCA system^18^, we first visualized the TAD (Topologically Associating Domain) and the distal enhancer-promoter interactions at the MYC gene locus (Extended Data Fig. 2). The median distance map generated by the ORCA system on a wide-field microscope showed a quantitatively similar profile to the one obtained on a previously reported ORCA system (Extended Data Fig. 2, 0.89 correlation coefficient) and showed good agreement with Hi-C data (Extended Data Fig. 2, 0.87 correlation coefficient)^22^.

To measure the chromatin structure and compaction at our engineered AAVS1 locus, we designed a DNA FISH probe library spanning our integrated reporter and the 150 kb region around it. Each set of readout probes in the library targets a 5 kb segment: thirteen contiguous segments (from -30 to 30 kb) around the reporter and three additional 5 kb segments upstream and downstream at 15 kb intervals (Fig. 1e). In addition, the entire region of interest is imaged in a second fluorescence channel in each round using fiducial probes as a reference for drift correction (Methods, ^23^). Median distances measured with these probe sets are reproducible across replicates with a correlation coefficient of 0.76 (Extended Data Fig. 3).

Since HEK293T cells possess three copies of chromosome 19 containing the AAVS1 locus, they may have different numbers of reporter integrations, ranging from 1 to 3. To increase the number of observations, we decided to use a clone that has multiple integrations. To ensure results are comparable between cell lines with different numbers of integrations, we isolated single clones that have either a single integration or multiple integrations (Extended Data Fig. 4a-c, Methods). KRAB recruitment resulted in full silencing and epigenetic memory formation in both single (A6) and double (B9) integration reporter lines (Extended Data Fig. 4d&e). As reactivation of either allele in the double integrant clone leads to an mCitrine-positive cell, the fraction of reactivated cells appears higher in the double integrant clone than the single integrant clone (Extended Data Fig. 4e); consistent with reactivation of a single allele, the fluorescence intensity of the reporter in the reactivated cells was lower than in cells that were never silenced for the double integrant clone (Extended Data Fig. 4f) but were the same in the single integrant clone (Extended Data Fig. 4g). These results confirm that the differences in apparent memory are due to the number of integrations, and stochastic gene reactivation of either one of the alleles (Extended Data Fig. 4h-j). Hereafter, the quantification of epigenetic memory was performed using the single integrant clone (A6), while chromatin structure analysis was conducted using the double integrant clone (B9). In addition, in order to avoid collecting data from AAVS1 loci without a reporter integrated, we filtered out traces without a positive readout signal at the integrated reporter.

### KRAB recruitment causes heritable chromatin compaction

After 5 days of KRAB recruitment, which resulted in gene silencing in nearly 100% of cells (Fig.1b middle), the median distance maps between chromatin segments measured by ORCA showed shorter distances compared to the no-dox cells (no recruitment, gene active) (Fig. 1f). The KRAB EEW25AAA mutant, which is incapable of silencing gene expression ^19^, served as a negative control and it did not induce chromatin compaction (Extended Data Fig. 5), suggesting that compaction is not caused by dox addition or rTetR binding to the TetO sites. Chromatin compaction was retained in irreversibly silenced cells even in the absence of KRAB recruitment, but not in reactivated cells (Fig. 1g), implying that chromatin compaction plays a role in irreversible gene silencing. Most pairwise segments that span across the reporter have shorter distances upon KRAB silencing and in irreversibly silenced cells after KRAB release compared to segments that do not contain the reporter between them (Fig. 1f,g bottom, Extended Data Fig. 6c), suggesting that chromatin compaction is localized around the reporter integration region (Extended Data Fig. 6d). To estimate local chromatin compaction from the single-cell traces, we measured the radius of gyration (Rg) around ±10 kb region at each position, and calculated the ratio of this radius of gyration upon KRAB recruitment (+dox) compared to no dox (Fig. 1h left, Methods). This local radius of gyration ratio showed a decrease at the tens of kb-scale around the reporter, especially downstream, suggesting that KRAB recruitment induces localized and asymmetric chromatin compaction (Fig. 1h).

It is of note that the tens of kb-scale chromatin structure changes observed here are much larger than the nucleation site, 9x TetO, which spans only 324 bp, consistent with spreading of chromatin modifications. Indeed, this observation aligns with asymmetric spreading of H3K9me3 upon KRAB recruitment for 5 days measured by CUT&RUN (Fig. 1i left,1j). However, most of the H3K9me3 enrichment was lost after KRAB release, not only in reactivated cells, but even in cells that remained irreversibly silenced for more than 30 days (Fig. 1i,j, Extended Data Fig. 1b). Both recently silenced (after 5 days) and irreversibly silenced cells lost active chromatin modifications H3K9ac and H3K27ac, consistent with complete gene silencing in these conditions (Fig. 1k,l, Extended Data Fig. 1c). The loss of H3K9me3 in irreversibly silenced cells prompted us to check for other modifications that are associated with chromatin compaction and epigenetic memory: H3K9me2 or H3K27me3^24^. However, neither of these appeared at the reporter integrated region in recently silenced or irreversibly silenced cells (Fig. 1m,n, Extended Data Fig. 1d). This result is consistent with the observation that inhibitors for these pathways do not reactivate the irreversibly silenced cells when used at concentrations known to attenuate gene silencing mediated by each modification^21,25^ (Extended Data Fig. 7). Instead, we found DNA methylation enrichment at the reporter in the irreversibly silenced population (cultured more than 30 days after release), but not at the end of KRAB silencing (5 days of recruitment) (Fig. 1o). In accordance with this observation, a DNA methyltransferase inhibitor could reactivate reporter expression in irreversibly silenced cells (Extended Data Fig. 7), suggesting that irreversible silencing may be maintained by DNA methylation. This is consistent with previous work showing that the combination of DNA methyltransferase inhibitor and deacetylase inhibitor reactivated randomly integrated reporter genes after KRAB-mediated silencing^7^. This finding suggests that de novo H3K9me3 may be involved in the initiation of compaction but is dispensable for the maintenance of compaction, and there is a handover from H3K9me3 to DNA methylation to establish epigenetic memory and maintain chromatin compaction.

### Single-cell chromatin structure is heterogeneous

Despite the clear difference in population-averaged distance maps, single chromatin traces are highly heterogeneous in both active and silenced cells (Fig. 1f,g top traces). It is not clear whether each chromatin structure comes from a distinct subpopulation in the set of possible conformations, or arises from a continuous spectrum of chromatin structures. To perform an unbiased analysis on this heterogeneous high-dimensional chromatin tracing dataset (78 dimensions for all combinations of pairwise segments within -30 kb to 30 kb), we first performed t-SNE dimensionality reduction and k-mean clustering with four kernels on aggregated data across all experimental conditions, then analyzed these clusters across individual conditions (Fig. 2a, Methods). The resulting clusters are not well separated in t-SNE space, suggesting a continuous distribution of chromatin states. Nevertheless, the distinct regions of this t-SNE space are associated with different levels of compaction: Clusters 1 and 2 display the most compacted and open conformations on average, and mainly occupy the negative side and the positive side of t-SNE1 axis, respectively (Fig. 2b). For each experimental condition, the projection of 3D chromatin configurations in this t-SNE space spreads broadly across all four clusters, and shows largely overlapping distributions with or without KRAB recruitment (Fig. 2c), suggesting that there is no obviously distinct subpopulation of chromatin conformations in the active state or upon silencing. The proportion of traces in the most compact conformation cluster (cluster 1) increases after gene silencing by KRAB recruitment for 5 days, but we also find a substantial number of traces with compact chromatin in cells where the gene is active (no dox) - in accordance with the broad distributions in the tSNE plot (Fig. 2d). Moreover, the proportion of traces in the most open conformation cluster (cluster 2) does not reach zero upon KRAB recruitment (Fig. 2e), even though nearly all cells are silent (Fig. 1b), indicating that the transition of chromatin conformation during silencing and epigenetic memory formation is not from fully open to fully compact, it is rather a moderate increase in the fraction of cells in the compact conformation.

**Fig. 2.**
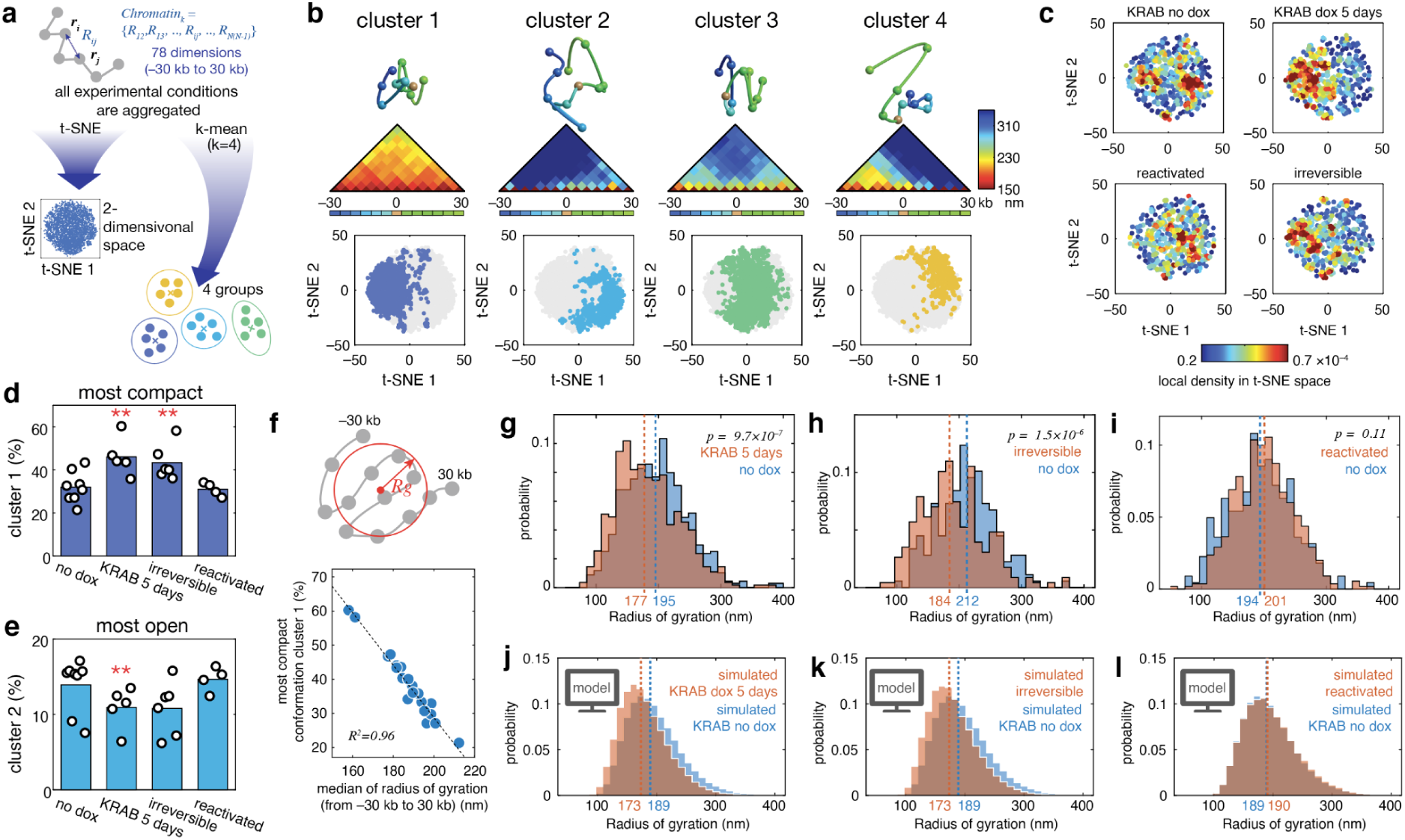
Single-cell chromatin structure is heterogeneous. (a) Schematic representation of single-cell analysis. Each chromatin trace has 78 dimensions. The aggregated data were mapped into 2-dimensional t-SNE space and clustered into four kernels using k-mean. (b) Representative 3D chromatin structures (top), median distance maps (middle) and territory in the t-SNE space of each cluster. (c) Distribution of t-SNE values from each experimental condition. The scatter plot is color-coded by local density. (d) Proportion of loci in cluster 1 (with the most compact conformation). Dots represent different biological replicates; each replicate quantified from 78 to 677 cells. ** indicate p<0.01 by Student’s *t*-test compared to no dox. Statistical comparison was made with only samples from the same replicate. (e) Proportion of loci in cluster 2 (with the most open conformation). (f) The proportion of loci in cluster 1 is plotted against the median of radius of gyration for each experimental condition. (g-i) Representative radii of gyration distributions of single-cell traces for (g) 5 days KRAB recruitment, (h) irreversibly silenced, or (i) reactivated conditions in red, and no dox control in blue. P values from a Wilcoxon Rank Sum test are shown. (j-l) Radii of gyration distributions from a 3D random-walk polymer simulation where the polymer step size was allowed to vary at each position to match the experimental median distance maps (Extended Data Fig. 9) for (j) 5 days of KRAB recruitment, (k) irreversibly silenced, or (l) reactivated conditions (red), versus simulations fitted to the no dox polymer (blue).

To see if the increase of compact conformations shown in the cluster analysis could be captured by a single parameter of the chromatin polymer, we calculated the radius of gyration of each single trace. Across different experimental conditions, the percentage of single-cell traces in the most compact chromatin cluster (cluster 1) correlates well with the median radius of gyration across all 3D chromatin traces (Fig. 2f), suggesting that the radius of gyration is sufficient to capture chromatin compaction in this experimental setup. Single-cell distributions of the radius of gyration at the end of KRAB recruitment and irreversibly silenced cells show a smaller median compared to the no dox one (Fig. 2g,h), while reactivated cells and no dox cells are not significantly different from each other (Fig. 2i). Despite the fact the median radius of gyration changes by ∼20 nm, which is a ∼10% reduction (see also Fig. 3e,f), there is a large overlap between the radii distributions of silenced populations and active ones (Fig. 2g,h) that comes from the high degree of heterogeneity of the single-cell polymer trajectories. To clarify how much of this heterogeneity could be the result of experimental noise, we developed a method to evaluate the degree of experimental noise. We assessed chromatin compaction before and after adding experimentally-derived noise to chromatin traces with clearly distinct configurations. We first measured the noise by re-hybridizing one of the segments (#7) at the end of chromatin tracing (Extended Data Fig. 8a). We measured the difference between the xyz coordinates of the first and second hybridizations, fit these data with a t-location scale distribution, and sampled random values from this distribution to simulate experimental noise (Extended Data Fig. 8b). We classified chromatin traces into either “t-SNE low” or “t-SNE high” based on the t-SNE 1 value, and then calculated their radius of gyration before and after adding noise to chromatin traces (Extended Data Fig. 8c). We found a clear segregation between t-SNE low and high populations even with simulated noise (Extended Data Fig. 8d), suggesting that our experimental noise could not explain the observed heterogeneity.

**Fig. 3.**
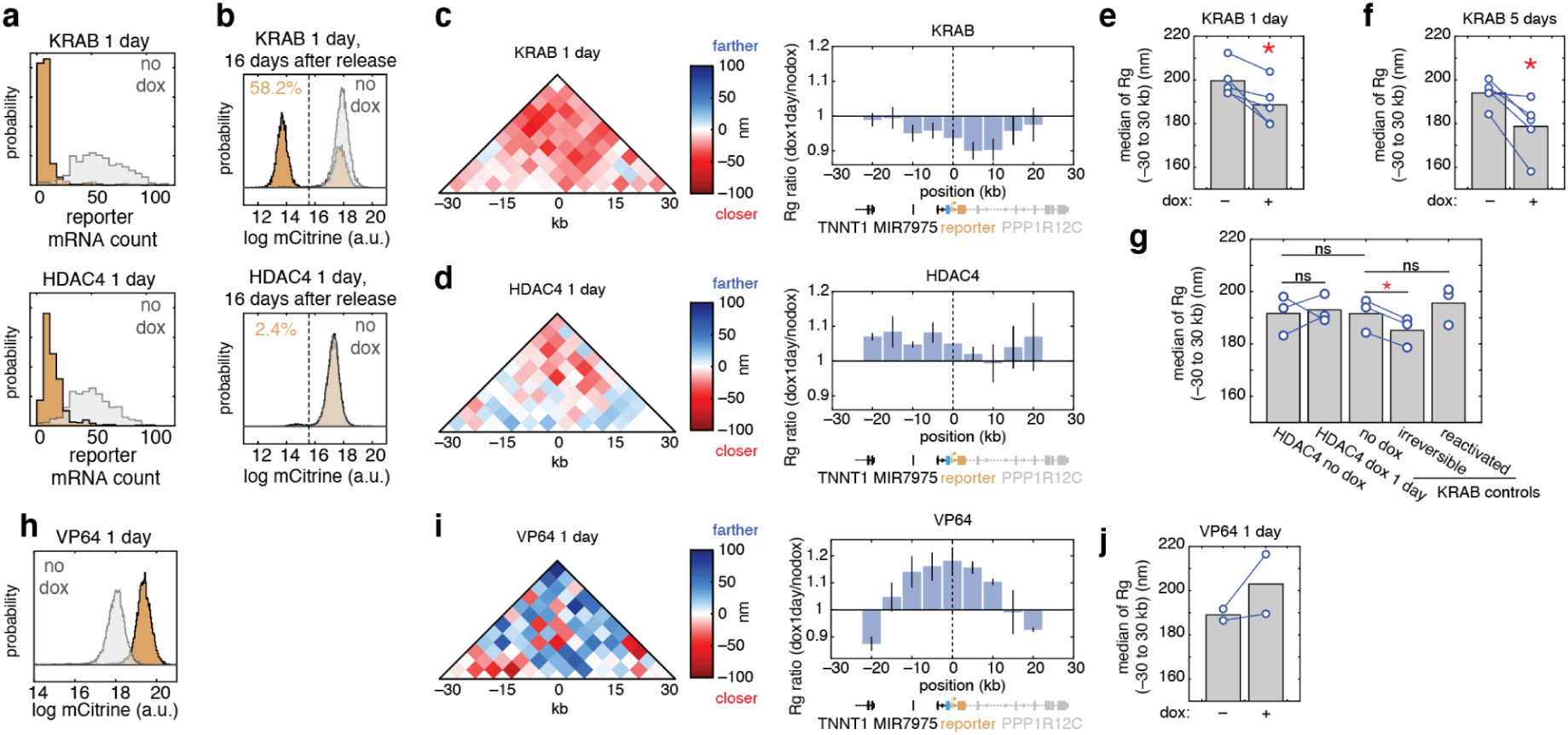
HDAC4 silencing does not lead to tens of kb-scale chromatin compaction. (a) Histograms show the distribution of reporter mRNA count measured at the single-cell level by RNA FISH (A6 clone). No dox control is shown in gray, and KRAB (top) or HDAC4 (bottom) recruitment by addition 1 µg/mL dox for 1 day is shown in yellow. (b) Histograms depict reporter mCitrine expression, measured by flow cytometry 16 days following KRAB (top) or HDAC4 (bottom) release (dox removal) (A6 clone). (c) (left) Subtracted median distance map between KRAB recruitment for 1 day versus no dox, and (right) local chromatin compaction measured by the local ratio of radius of gyration (±10 kb around each position) compared to no dox. Bars show mean ± s.e.m. (d) Same as C, for HDAC4 recruitment. (e-f) Median of the radius of gyration (from -30 to 30 kb) with or without KRAB recruitment for (e) 1 day or (f) 5 days. Each blue line connects data from the same replicate (imaged at the same time on the same slide). * indicate p<0.05 by Student’s *t*-test compared to no dox. (g) Median of the radius of gyration (from -30 to 30 kb) for HDAC4 1 day recruitment cells and no dox, and for indicated KRAB controls imaged at the same time. Blue lines as in e,f. * indicate p<0.05 by Student’s *t*-test. (h) Histogram of reporter mCitrine expression as measured by flow cytometry upon VP64 recruitment by addition 1 µg/mL dox for 1 day (A6 clone). (i) Same as c and d, for VP64 recruitment. (j) Same as e and f, for VP64 recruitment.

We next hypothesized that the observed heterogeneity could arise from the dynamic polymer nature of the chromatin fiber that samples many conformations. In fact, the dynamic nature of chromosome folding has been proposed before to explain the lack of distinct clusters of 3D chromosome conformations observed in single-cell data for CTCF-mediated chromatin looping^26^ and long-range enhancer-promoter interactions^27^. To analyze the extent of heterogeneity expected from a flexible fiber, we developed a simple 3D random walk polymer model (Methods), and analyzed the predicted single-cell heterogeneity. In this model, we sampled step sizes between 70 to 210 nm, and used Bayesian optimization to find the step sizes that match our experimentally-measured median distance maps (Extended Data Fig. 9, Methods). The radii of gyration distributions show a large overlap between polymers fitted to distance maps from silenced cells and those fitted to no dox conditions (Fig. 2j-l), similar to the experimental data (Fig. 2g-i). These results indicate that the heterogeneity of chromatin structure could arise from the flexible polymer nature of chromatin. In summary, even under stable active gene expression, chromatin displays compact conformations without distinct subpopulations, and it becomes more biased, but not completely shifted, towards compact conformations upon silencing by KRAB recruitment.

### Transcriptional silencing does not require or produce chromatin compaction at the tens of kilobases scale

To gain a deeper understanding of the relationship between chromatin compaction and gene expression, we tested other perturbations. As an alternative silencing perturbation, we used histone deacetylase HDAC4, which is known to remove acetyl groups from histones^28^ and repress gene expression^15^. Both KRAB and HDAC4 silenced reporter gene expression in almost all cells within 1 day of recruitment, as quantified by RNA FISH (Fig. 3a). However, in contrast to KRAB recruitment, HDAC4 recruitment did not exhibit epigenetic memory (Fig. 3b), and did not lead to the clear chromatin compaction that is observed in KRAB recruitment at the tens of kb scale (Fig. 3c,d). We tested if there is significant chromatin compaction, as measured by the radius of gyration, in these samples. As expected, we observed a significantly smaller Rg after both 1 day or 5 days of KRAB recruitment (Fig. 3e,f). In contrast, we did not find a significant reduction in the Rg after HDAC4 recruitment compared to no dox (Fig. 3g). For positive and negative controls imaged at the same time on the same slide, we used the KRAB-containing cells, and indeed we detect a significant reduction in Rg between KRAB irreversibly silenced cells and KRAB no dox, but no significant difference between KRAB and HDAC4 cell lines in the absence of dox (Fig. 3g). This finding suggests that transcriptional silencing does not require chromatin compaction at this length scale (10-20 kb), and that transcriptional silencing by itself does not cause the type of chromatin compaction over 10-20 kb we observed with KRAB recruitment. However, even with HDAC4 recruitment we cannot exclude the possibility of weak chromatin compaction that is under the detection limit of this experimental setup or chromatin compaction at a smaller scale (under 5 kb, our step size).

As an additional perturbation to transcription, we tested recruitment of VP64, which is known to activate gene expression. VP64 recruitment increased reporter gene expression compared to the baseline no dox expression (Fig. 3h), and also resulted in a more open chromatin conformation (Fig. 3i,j), in line with previous observation of chromatin decondensation upon VP64 recruitment^29^. This increase in open chromatin spans approximately 20 kb (Fig. 3i), a similar length scale to chromatin compaction upon KRAB recruitment (Fig. 3c), suggesting that active chromatin states can also spread as previously observed^30^.

### Chromatin compaction at the end of silencing predicts epigenetic memory

Since we measured greater compaction upon silencing followed by memory (by KRAB), as opposed to silencing without memory (by HDAC4) (Fig. 3c,d), we set out to systematically investigate the potential association between chromatin compaction and epigenetic memory. We modulated the level of epigenetic memory by varying the duration recruitment and by using a KRAB mutant with reduced memory. This KRAB mutant, which we identified in a previous functional screening assay with K562 cells^19^, contains a point mutation (Y46A) in the A-box domain that binds to its main corepressor KAP1^31^. We silenced reporter expression with KRAB or KRAB Y46A for 1 day or 5 days. At the end of recruitment, double integrant cells were fixed for chromatin tracing, and single integrant cells were cultured in dox-free media to measure epigenetic memory (Fig. 4a). Upon recruitment, regardless of the point mutation or the duration of recruitment, more than 90% of cells across all conditions are silenced as confirmed by RNA FISH (Fig. 4b, top). However, despite similar silencing profiles, we found different levels of epigenetic memory after dox removal, ranging from 20% to 80% across treatments (Fig. 4b, bottom), with 5 days recruitment of WT KRAB having the highest percent of irreversibly silenced cells and 1 day Y46A KRAB the lowest. Thus, the mutant KRAB or shorter recruitment both decrease epigenetic memory measured 16 days after release, rather than gene silencing at the end of recruitment. Despite a similar percentage of silenced cells, chromatin tracing revealed less compaction at the end of KRAB Y46A mutant recruitment compared to WT KRAB (Fig. 4c). Moreover, the radius of gyration of the single-cell chromatin traces at the end of recruitment shows a graded reduction from ∼200 nm to ∼180 nm directly proportional with the increase in epigenetic memory measured 16 days later (Fig. 4d, Extended Data Fig. 10). The no silencing controls, including KRAB EEW25AAA mutants, consistently show a radius of ∼200 nm (Fig. 4e). These findings indicate that chromatin compaction at the end of silencing is predictive of epigenetic memory weeks later.

**Fig. 4.**
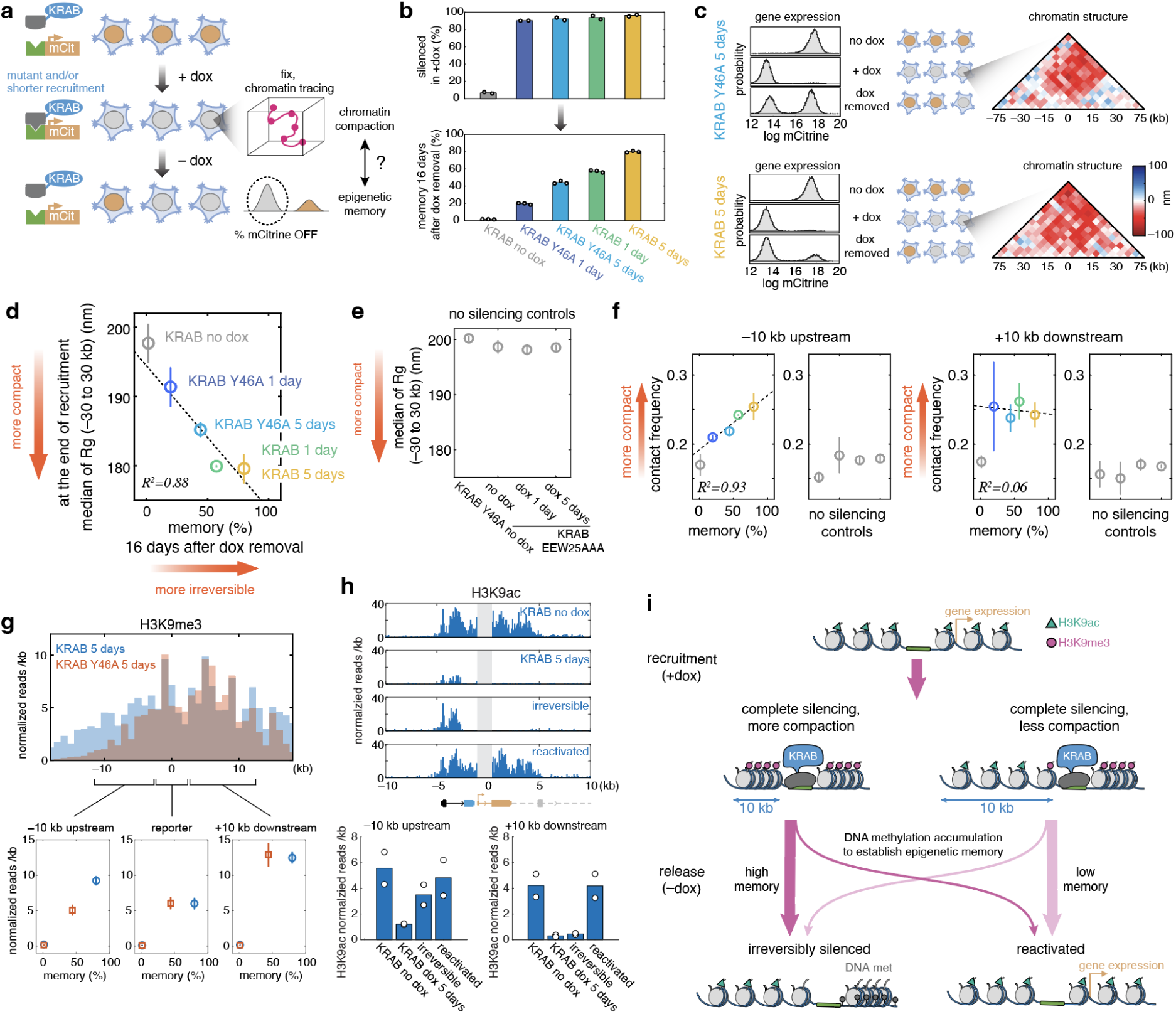
Chromatin compaction predicts epigenetic memory. (a) Schematic representation of experimental workflow for quantifying epigenetic memory and chromatin compaction. Epigenetic memory was defined as the percentage of irreversibly silenced cells after dox removal. (b) Silencing and epigenetic memory (as percentage of cells with mCitrine off) across varying strengths of KRAB recruitment tuned by Y46A mutation and shorter recruitment duration. Percentage of silenced cells was quantified by (top) RNA FISH at the end of recruitment and (bottom) flow cytometry for epigenetic memory 16 days after releasing KRAB. (c) Representative flow distributions (no dox, +dox 5 days, and dox removed 16 days) and subtracted median distance maps measured by ORCA at the end of recruitment (+dox 5 days) for (top) KRAB Y46A or (bottom) KRAB WT. (d) Median radius of gyration for the -30 to 30 kb region around the reporter at the end of recruitment plotted against epigenetic memory (percentage of cells off 16 days after dox removal). Circles show the average of two replicates, vertical bars show maximum and minimum values. Dashed lines show linear regressions across conditions after excluding the no dox at memory % = 0. (e) Median radius of gyration in no silencing control conditions (see also Extended Data Fig. 5 for KRAB EEW25AAA). Circles show the average of two replicates, vertical bars show maximum and minimum values. (f) Contact frequency (fraction of detected readout signals within 150 nm from the reporter) between reporter and (left) -10 kb upstream or (right) +10 kb downstream segments plotted against different memory levels (% cells silent as in d). The Colors are the same as in d. The conditions for no silencing controls are the same as in e. Circles are average of two replicates, vertical bars show maximum and minimum values. Dashed lines show linear regressions across conditions after excluding the no dox at memory % = 0. (g) (top) Genome traces showing the normalized number of reads after CUT&RUN against H3K9me3 as a function of distance around the reporter integration site (at 0 kb), upon WT KRAB recruitment (blue) or KRAB Y46A recruitment for 5 days (orange). (bottom) CUT&RUN data from two replicates are averaged. Quantification of the normalized H3K9me3 CUT&RUN reads within (left) the 10 kb upstream region, (middle) the reporter integrated region, or (right) the 10 kb downstream region under no dox (0 memory points) or dox for 5 days to recruit KRAB (blue) or KRAB Y46A (orange) plotted against epigenetic memory. Bar shows maximum and minimum values from two replicates. (h) (top) Genome traces showing the normalized number of reads (Averaged from two replicates) after CUT&RUN against H3K9ac under no dox, 5 days of KRAB recruitment, irreversibly silenced, or reactivated conditions. The human EF1alpha promoter driving the reporter gene is masked with gray rectangles since sequencing reads also align to the endogenous copies of EF1alpha that are not targeted with KRAB and are therefore expected to retain acetylation. (bottom) Quantification of the normalized H3K9ac at (left) the 10 kb upstream region, (middle) the reporter integrated region excluding EF1alpha promoter, (right) 10 kb downstream region. (i) Schematic of chromatin state transitions upon KRAB recruitment.

In order to elucidate which region of the compacted chromatin contributes to the prediction of the epigenetic memory, we calculated contact frequency between the reporter and other segments. We found that the region that is 10 kb upstream shows a graded increase in contact frequency to the reporter that is correlated with epigenetic memory, while the 10 kb downstream region showed a consistently high contact frequency across all silencing conditions (Fig. 4f). This finding suggests that the downstream region is highly compacted immediately upon silencing, while the upstream region compacts as cells become more likely to be epigenetically silenced. The contact frequency between the upstream segments and the reporter shows higher predictability overall, defined as the R^2^ value after linear regression (median of R^2^ = 0.57), compared to the downstream segments (median of R^2^ = 0.14) (Extended Data Fig. 11a). This observation is also confirmed by other metrics such as average distance of upstream versus downstream segments from the reporter, or the number of segments included within a 150 nm radius from the reporter (Extended Data Fig. 11b-e). Therefore, in this reporter system, compaction of the upstream region after silencing is more predictive for future epigenetic memory.

This asymmetric chromatin compaction and correlation with memory is in line with the H3K9me3 profile across tens of kb around the reporter: while the lower memory KRAB Y46A induces de novo H3K9me3 to a similar level with wild-type KRAB at the reporter and the downstream region spanning 10 kb, H3K9me3 enrichment at the upstream region is lower than wild-type KRAB (Fig. 4g). We hypothesized that histone acetylation upstream of the reporter, around the actively transcribing PPP1R12C gene, can oppose the spreading of histone methylation, since H3K9ac needs to be removed before H3K9me3 deposition. We found a wide H3K9ac distribution across the reporter integration site in the absence of KRAB recruitment (no dox) and reactivated cells. We noticed a region of H3K9ac enrichment in the upstream region (around the PPP1R12C promoter and its first intron) that is more resistant to KRAB-mediated silencing. Acetylation in this upstream region decreases after 5 days of recruitment and reappears in irreversibly silenced cells, while H3K9ac at the downstream region is irreversibly lost (Fig. 4h). These results suggest that the spreading of histone methylation and chromatin compaction beyond the recruitment site depend on the presence of active modifications in the region.

These results lead to the following model of how chromatin states transition during silencing and epigenetic memory formation (Fig. 4i): Epigenetic perturbation by KRAB recruitment can either induce a wide H3K9me3 domain and chromatin compaction across 10-20 kb that is likely to be converted to DNA methylation and result in irreversible silencing after KRAB release (left), or a smaller H3K9me3 domain and less chromatin compaction that is more likely to be lost (converted to active modifications such as histone acetylation) and result in gene reactivation (right). This model raises the hypothesis that the level of chromatin compaction achieved at the end of recruitment across the tens of kb-scale, beyond the targeted gene itself, affects epigenetic memory establishment.

### Chromatin compaction upon differentiation correlates with irreversible fate commitment in mESCs

We wanted to see if this predictive relationship between chromatin compaction and irreversible gene silencing plays an important role in a biological process where genes are epigenetically silenced, for example during cell differentiation. H3K9me3 is known to play a role in maintaining cellular identity through irreversible gene repression^24^. A recent study has demonstrated that H3K9me3 enrichment at the Nanog gene locus, a key factor conferring pluripotency, is important for timing irreversible fate commitment during mouse ES cells (mESCs) differentiation^32^. In order to modulate and assess the percentage of cells that are irreversibly committed, we initially grew cells in differentiation media for varying time periods, and then replated 500 cells in stem cell media, allowing the ones that are not irreversibly committed to form ES colonies^32^ (Fig. 5a). Cell morphology clearly changed upon differentiation (Fig. 5b), and we confirmed previously reported changes in gene expression of cell marker genes^32,33^ by qPCR (Extended Data Fig. 12). We observed an increase over time in irreversible commitment to differentiated cells that cannot revert to mESCs, as evidenced by the decreasing number of cell colonies after replating (Fig. 5b insets, 5c), consistent with previous findings^32^. While the Nanog gene was silenced as early as day 1 (Fig. 5d), the chromatin structure around the Nanog locus exhibited gradual compaction over a longer time period (as observed with KRAB), with compaction increasing especially at day 3 (Fig. 5e). Similarly to our KRAB-targeted reporter, this chromatin compaction could be measured as a decrease in the median radius of gyration, and it correlated well with the decrease in the number of colonies (i.e. increase of cell fate memory) (Fig. 5f). For this particular locus, we found that the contact frequency between the Nanog promoter and the downstream region is more predictive than the upstream region (Fig. 5g, Extended Data Fig. 13a). In this case, the downstream region overlaps the neighboring Slc2a3 - an active gene enriched in histone acetylation (Extended Data Fig. 13b,c), while in the case of the KRAB-targeted reporter, the region with active modifications is upstream. Taken together, these results suggest that chromatin compaction across tens of kbs, and especially across nearby regions containing active regulatory elements, can be a better predictor of irreversible fate commitment than the level of gene expression.

**Fig. 5.**
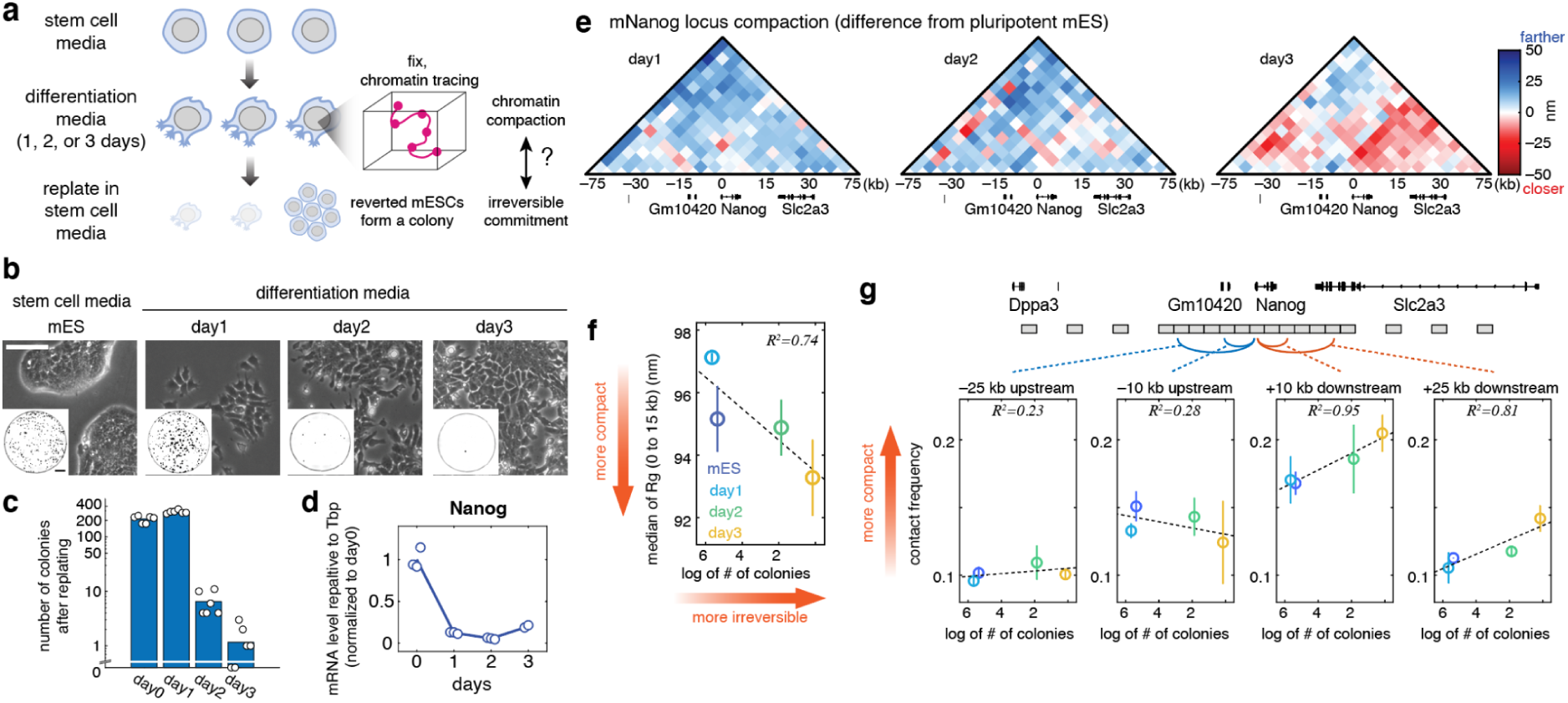
Chromatin compaction correlates with irreversible fate commitment in mouse ES cell differentiation. (a) Schematic representation of the experimental workflow for quantifying irreversible fate commitment and chromatin compaction during mESCs differentiation in N2B27 media. (b) Example images of mouse embryonic stem cells before differentiation and at different time points of differentiation. Scale bar (white): 100 µm. Insets show colonies after replating differentiated mES cells in stem cell media. Scale bar for insets (black): 250 mm. (c) Number of colonies after replating, calculated from insets in Fig. 5a. (d) Nanog expression level was measured by RT-qPCR. The mRNA levels were quantified relative to Tbp, then normalized to the average mRNA level at day 0. Line plots show the average across two or three biological replicates (dots). (e) Subtracted median distance maps upon differentiation compared to pluripotent mES cells. Data from two replicates are averaged. (f) Median radius of gyration of the 0 to 15 kb region is plotted against the log of the number of colonies in Fig. 5b. Circles show the average of two replicates, vertical bars show maximum and minimum values. (g) Contact frequency (fraction of detected readout signals within 100 nm from the Nanog promoter) between the Nanog promoter and -25 kb, -10 kb upstream, or +0 kb, +25 kb downstream segments. Circles show the average of two replicates, vertical bars show maximum and minimum values. Dashed lines show linear regression.

## Discussion

Here, we measured how the 3D chromatin structure changes in single cells at a locus where we finely control gene silencing and the level of epigenetic memory by recruiting different repressors for different periods of time. This system allowed us to quantify changes in the chromatin states that are associated with epigenetic memory establishment separately from gene silencing. We show that repression associated with epigenetic memory leads to chromatin compaction across tens of kbs around the target gene, and the level of compaction quantitatively predicts the fraction of cells with epigenetic memory. We also show that chromatin compaction quantitatively predicts irreversible epigenetic fate commitment during mouse ES cells differentiation. On the other hand, we do not observe this long-range chromatin compaction in cells that are silenced reversibly, suggesting that gene silencing alone is not sufficient to cause compaction at this length-scale and that compaction is not necessary for silencing, but rather for epigenetic memory. Moreover, we show that in irreversibly silenced cells compaction is consistently maintained for weeks after the end of silencing into the memory phase. However, the molecular underpinnings of the compaction associated with memory change over time: compaction at the end of silencing correlates with H3K9me3 spreading across the locus, while in irreversibly silenced cells (two weeks after repressor release) H3K9me3 modifications are lost from the locus and instead we find DNA methylation associated with chromatin compaction (Fig. 1).

Our results suggest that in our system histone methylation is converted into DNA methylation as epigenetic memory is established and maintained. Indeed, crosstalk between H3K9me3 and DNA methylation has been reported^34^, and DNA methylation often silences gene expression irreversibly^7,15^. It is also known that DNA methylation accumulates at a gene after long continuous recruitment of KRAB^35^ or HP1^8^ in mouse embryonic stem cells. Our observation that DNA methylation enriches mostly after the release of KRAB domain suggests that the process is mediated by an endogenous system that converts H3K9me3 to DNA methylation, not necessarily by co-factors associated with the KRAB domain. Alternatively, a recent study suggested that MPP8, initially recruited by H3K9me3, maintains the silencing of transposable element LINE1 even in the absence of H3K9me3^36^. These results imply that despite the existence of different molecular modules that can maintain epigenetic memory, and the fact that there can be handover from one type of module to another during the memory phase, the common signature across these modules is chromatin compaction at the tens of kb scale.

What is the role of this tens of kb-scale chromatin compaction in the irreversibly silenced state? One possibility is that it is making DNA inaccessible to transcription factors. However, DNA methylation alone can suppress gene expression by directly inhibiting transcription factor binding to the genome^37^. Another possibility is that compaction contributes to the maintenance of epigenetic modifications. Epigenetic modifications spread across the genome^8,21,38^ by either chromatin looping, reader-writer feedback, or diffusion of locally concentrated histone modifiers^8,39,40^. Increasing compaction, and hence chromatin contact frequency, would therefore help propagation or maintenance of epigenetic modifications^41–43^.

How does the level of chromatin compaction we observe translate to chromatin density and how does it compare with other heterochromatin? In conditions that lead to high epigenetic memory, changes in the radius of gyration range from 0.8x to 0.9x, corresponding to reductions in chromatin volume by almost a half or a quarter (since 0.8^3^ = 0.512 and 0.93^3^ = 0.729) and therefore a big increase in nucleosome density. This magnitude is similar to direct measurements of nucleosome density using electron microscopy-based method, ChromEMT, which revealed twice the density of heterochromatin compared to euchromatin^44^.

Despite the strong compaction observed in the population average upon KRAB silencing and memory, at the single-cell level we find that chromatin structure is heterogeneous, with open and compact conformations present in both active and silenced cells. This heterogeneity suggests that even with H3K9me3 enrichment across more than 10 kb, chromatin does not undergo an all-or-none transition from fully open to fully compact, but instead it is dynamic in both the active and silenced states. Similar single-cell results were recently reported for Polycomb repression in mouse ES cells^43^. It is possible that the change in average compaction is a result of different transition rates between open and compact conformations in active versus silent loci. These dynamics could be directly measured using live imaging of chromatin, as has been done for enhancer-promoter contacts^45^ or CTCF- and cohesin-mediated chromatin looping^46^.

We show that in order to predict the level of epigenetic memory of a gene, we need to measure chromatin compaction at a larger region that extends beyond that gene. Our chromatin tracing data at 5 kb resolution allowed us to perform a more detailed analysis as to which segment contributes most to epigenetic memory as opposed to silencing. In both the synthetic reporter and the Nanog locus, the target gene compacts soon upon silencing, but compaction of active regions beyond the target gene are more important for epigenetic memory. In the case of our AAVS1 integrated reporter, the active region important for memory was situated upstream, and showed histone acetylation enrichment around the promoter and first intron of the PPP1R12C gene; in the case of Nanog, the active region that compacts upon irreversible fate commitment was situated downstream and showed histone acetylation enrichment around the promoter of the Slc2a3 active gene. In theoretical models of epigenetic regulation, the antagonism between histone acetylation and methylation is hypothesized to be an important parameter for setting the extent of epigenetic modifications spreading and their spatial patterns^40,47^. Our results indicate that this type of antagonism between active and repressive modifications could play a role in epigenetic memory formation as well.

We also show the quantitative link between chromatin compaction and irreversible fate commitment holds in a very different biological process, during cell differentiation: chromatin compaction at the Nanog locus during mESC differentiation correlates with the number of cells that are irreversibility committed to the differentiated state and cannot revert back to form mES cell colonies after replating them in stem cell media. Changes in local chromatin accessibility associated with irreversible fate transitions during differentiation have been reported in other systems: In a granulocytic differentiation model derived from murine bone marrow, irreversible fate commitment is associated with irreversible loss of chromatin accessibility measured by ATAC-seq^48^. Moreover, genome-wide chromatin accessibility and gene expression analyses of hair follicle differentiation in mouse skin have shown that local chromatin accessibility changes prior to gene expression and foreshadows lineage choice^49^. In fruit fly eye development, chromatin decompaction at the *ss* gene locus (imaged by DNA FISH) precedes R7 subtype differentiation^50^. These studies suggest that chromatin structure at the end of an epigenetic perturbation or differentiation stimulus may determine the potential of gene expression in future. Our results showing a quantitative correlation between initial chromatin compaction upon direct epigenetic perturbations and the fraction of cells with irreversible gene silencing at a later time suggests that chromatin architecture at the tens of kb scale could be used to predict cell fate in other biological scenarios where epigenetic silencing comes into play. Genome-wide analyses of chromatin architecture, such as Hi-C or multiplexed DNA FISH could be used to test predictions of irreversible fate commitment during differentiation to different cell types (before the end of the differentiation protocol) or during other processes associated with long-term epigenetic modifications, such as aging^51^ or inflammation^52^.

## Materials and Methods

### Human cell culture and reporter cell line generation

Human embryonic kidney cells (HEK293T, Takara Bio #632180)) were cultured in DMEM-GlutaMAX media (10566024, Gibco) supplemented with 1x Penicillin-Streptomycin-Glutamine (10378016, Gibco) and 10 % FBS (FB-15, Omega Scientific). The reporter cell line was generated in previous work^55^, by co-transfecting cells with TALEN-L (Addgene #35431), TALEN-R (Addgene #35432) and pJT039 (AAVS1-9xTetO-pEF-IGKleader-Cas9site-hIgG1_FC-Myc-PDGFRb-T2A-Citrine-PolyA) reporter donor (Addgene #161927)^19^, followed by antibiotic selection with 0.25 µg/mL puromycin. To isolate single clones, cells were diluted at 100 cells/20 mL, and then dispensed into a 96-well plate at 200 µL per well (∼1 cell/well). After 12 days of culture, wells with a single colony were selected and further expanded. We measured mCitrine reporter expression levels using flow cytometry ZE5 (Bio-Rad), and extracted genomic DNA with QuickExtract DNA Extraction Solution (QE09050, Lucigen). PCR on the genomic DNA was performed to identify the presence of chromosomes with or without reporter integration^56^. To verify the number of integrations, we extracted genomic DNA from 2 million cells using DNeasy Blood & Tissue Kit (69504, Qiagen) and performed droplet digital PCR (ddPCR) through a service provided by MOgene (Extended Data Fig. 4b). Note that the ddPCR result is not always concordant with the results of the genomic PCR: for example, the B9 clone did not show a positive band for PCR on the wild-type chromosome, suggesting all three copies of chromosome 19 have the reporter, whereas ddPCR suggested the B9 clone has two integrations instead of three. As an alternative way to check the integration, we performed DNA FISH using ORCA probes (described in ORCA: probe design and synthesis section in detail), and calculated the percentage of loci with fiducial signal (detected across all chromosomes 19 in a cell with ∼3000 cy3 probes across the entire imaged region) that have a positive readout DNA FISH signal at the reporter (that detects the integrated reporter segment with lower efficiency since it only uses ∼150 cy5 probes). The percentage of reporter-positive chromosomes (Extended Data Fig. 4c) quantitatively matches the ddPCR results (Extended Data Fig. 4b), therefore we concluded A6 has a single integration, B9, B11 and D5 have two, and F5 has three integrations. One possible cause for the lack of signal in the genomic PCR for some of the double integrant clones could be the formation of indels that abolish primer binding near the TALEN cut site.

The wild-type KRAB domain from ZNF10, KRAB Y46A mutant, KRAB EEW25AAA mutant, and VP64 were individually cloned as fusions to rTetR(SE-G72P)^20^ using the backbone from pJT126 lenti pEF-rTetR(SE-G72P)-3XFLAG-LibCloneSite-T2A-mCherry-BSD-WPRE (Addgene #161926) digested with Esp3I-HF. They were delivered by lentivirus^19^. Wild-type KRAB, KRAB Y46A mutant, KRAB EEW25AA mutant, and VP64 positive cells were selected by 10 µg/mL blasticidin S (A1113903, Gibco) and sorted by gating for mCherry-positive signal.

The pPB:PGK-H2B-mIFP-T2A-rTetR-DNMT3B1-polyA-zeo plasmid was transfected in the HEK293T reporter cell lines together with Super PiggyBac Transposase to obtain cells stably expressing rTetR-DNMT3B. Cells were selected in 60 µg/mL zeocin (ant-zn-05, InvivoGen) followed by cell sorting by gating for mIFP-positive signal. Cell sorting was performed twice to enrich mIFP positive cell population.

The rTetR(SE-G72P)-HDAC4-T2A-mTurquoise plasmid was generated by PCR amplifying full length human HDAC4 with appended gibson homology arms from pLB37_PB_pGK-H2B-mIFP-T2A-rTetR-HDAC4-zeo (Addgene #179440) using Q5 hot start polymerase (NEB M0494S). A lentiviral backbone containing pEF1alpha-driven expression of 3xFLAG-tagged rTetR(SE-G72P) was then linearized by Esp3i digestion and the HDAC4 PCR fragment was annealed downstream into the open reading frame with gibson assembly using NEB HiFi mastermix (NEB 2621L). After delivering the construct by lentivirus, HDAC4-positive cells were sorted by gating for mTurquoise-positive.

To sort reactivated and irreversibly silenced cells after the KRAB recruitment and release, double reporter integrant cells (B9) were treated with 1 µg/mL doxycycline (dox; 40902, Tocris) for 5 days and then cultured without dox for 22 days. A population where both copies of the reporter reactivated (top ∼2% in fluorescence intensity) and a completely silenced population were sorted using the SH800 cell sorter (SONY).

For the validation of ORCA on a wide-field microscope, hTERT-RPE1 cells were used. The cells were cultured in DMEM-GlutaMAX media supplemented with 100 U/mL Penicillin-Streptomycin (15140122, Gibco) and 10 % FBS. The cells carry a synthetic reporter gene but it was not utilized in this research.

### Epigenetic modifications inhibitor treatments

To inhibit H3K27 methyltransferase, H3K9 methyltransferase, or DNA methyltransferase activity, cells were treated with Tazemetostat (S7128, Selleck Chemicals), Chaetocin (13156, Cayman Chemical Company), or 5-Aza-2’-deoxycytidine (A3656-5MG, Sigma-Aldrich) respectively for 4 days at concentrations indicated in Extended Data Fig. 7. To assess the stability of reporter expression after drug removal, cells were washed with 250 µL of DPBS twice and cultured for 8 days.

### Flow cytometry

To silence reporter expression and measure epigenetic memory, cells were cultured in the presence of 1 µg/mL doxycycline (dox; 40902, Tocris). During the dox-treatment phase, culture medium was exchanged every day. For removing dox, cells were spun down and washed with 250 µL DPBS (25-508, Genesee Scientific) twice, then resuspended in culture media with no dox. Cells were passaged every 2-4 days. Before measuring fluorescence on flow cytometry, cells were resuspended in 1X HBSS (14175095, Gibco) supplemented with 1 mM EDTA (BM-711, Boston BioProducts) and 0.5 mg/mL BSA (22014, BIOTIUM). Flow cytometry was performed to measure mCitrine, mCherry or mTurquoise fluorescence intensity on ZE5 (Bio-Rad). Flow cytometry data were gated by forward and side scatter to select live single cells, as well as on mCherry or mTurquoise positive signal to select cells expressing the rTetR-fused chromatin regulators. To calculate the percentage of cells with epigenetic memory, cells with the mCitrine fluorescence level lower than 15.5 in log scale were classified as silenced. All analyses were performed in MATLAB using Easyflow developed by Dr. Yaron Antebi (https://github.com/AntebiLab/easyflow).

### ORCA: probe design and synthesis

We designed ORCA probe sets as previously described ^18,23^. In brief, the barcoded oligos were designed on the sense strand to avoid the probes hybridizing onto mRNA. The algorithm selects probes without homology to human repetitive sequences, and with GC contents within the range of 20 to 80%. Oligos that can hybridize to each other were also removed. The code is available at Alistair Boettiger lab’s Github repository (https://github.com/BoettigerLab/ORCA-public). The probes were ordered from GenScript, and amplified as previously described ^23^. Briefly, the primary probes were amplified using Phusion® High-Fidelity PCR Master Mix with HF Buffer (M0531S, NEB) by monitoring the amplification on a qPCR machine (CFX Connect, Bio-Rad) to make sure amplification is in the linear range. Single-stranded RNA was synthesized from the PCR product using Hiscribe T7 Quick High Yield RNA Synthesis Kit (E2050S, NEB). The ssRNA was then converted into ssDNA using Maxima H Minus Reverse Transcriptase (EP0753, Thermo Scientific), and used as primary probes. The concentration of primary probes was measured using the Qubit ssDNA Assay Kit (Q10212, Thermo Scientific), and it usually ranged between 80 to 180 ng/µL.

### ORCA: cell culture on a coverslip

To minimize batch effects, we always plated dox treated cells and no dox control cells on the same coverslip. To create isolated cultures on a coverslip, we made a PDMS multi-well device on top of the imaging slide as follows: Silicone Elastomer Base and Curing Agent (Sylgard 184; 4019862, Dow Corning) were mixed at 10:1 ratio and poured into a 10 cm dish. The dish was placed in a desiccator to degas the PDMS for 1 hour. The degassed PDMS was then left on a bench at room temperature to cure. We used the PDMS at least 1 week after curing to get electrochemically stabilized PDMS. A 3.17 cm(1.25 inch)-by-1.9 cm(0.75 inch) PDMS chunk was cut out from the cured PDMS, then 13 of 3 mm holes were created using a disposable biopsy punch (MedBlades). The PDMS multi-well was pressed onto a 40 mm coverslip (Bioptechs). To plate HEK293T cells, the bottom of each PDMS well was coated with 25 µL of 5% Matrigel (356231, Corning) in DMEM-GlutaMAX + 1x PenStrepGlutamate + 10% FBS in a 37°C tissue culture incubator for one hour after brief centrifugation to spin down the marigel solution to the bottom of the wells in a 6-cm dish. Meanwhile, cells were trypsinized with 0.25% trypsin-EDTA (252-000-56, Fisher Scientific) for 5 minutes, spun down and resuspended at the density of 8000 cells / 25 µL with or without doxycycline. After removing the matrigel solution, 25 µL of the resuspended cells were loaded into each well and spun down at 700xg for 1 minute at room temperature. A piece of Kimwipes (34155, Kimberly-Clark) wetted with 2 mL of water was placed in the 6 cm dish to humidify the dish. After confirming cells are at the bottom of wells, the coverslip in a 6 cm dish was then incubated in the TC incubator for 24 hours. After fixing the cells the following day (detailed in the next section), the PDMS multi-well device was removed, washed with 70% ethanol and Milli Q water, and air-dried to reuse in future experiments. For experiments involving more than 1 day of dox treatment, cells were cultured in the presence of dox in a culture dish before plating on the coverslip.

To plate hTERT-RPE1 adherent cells, a 40 mm coverslip was pre-coated with Cellmatrix Type I -P (Collagen, Type I, 3 mg/mL, pH 3.0) (634-00663, Fujifilm) diluted in DPBS for 1 hour in the culture incubator.

### ORCA: primary probe hybridization

We performed primary probe hybridization for ORCA experiments as previously described with some modifications ^18,23^. Cells were cultured in 25 µL of the culture media in the PDMS multi-well device, and fixed by adding 25 µL of 8% formaldehyde (15714-S, Electron Microscopy Sciences) in 1xPBS (f.c. 4%) for 10 minutes. The formaldehyde solution was then aspirated, and the PDMS multi-well device was carefully removed. Cells were further fixed with 2 to 4 mL 4% formaldehyde for 10 minutes by completely submerging the coverslip in the formaldehyde solution followed by 3 washes with 1xPBS (AM9625, Invitrogen). Hereafter, all treatments and washes were performed using 2 to 4 mL of each solution to thoroughly treat the coverslip unless otherwise noted. We also want to note that the solutions should not be poured directly onto the fixed cells, to avoid disrupting the cells. For long-term storage of the fixed cells, the coverslip was washed with 70% ethanol once, and submerged in 70% ethanol, sealed with parafilm, and stored at -20°C in a paper box. Before the following steps were performed, the coverslip was placed at room temperature for 10 minutes and washed with 1xPBS. Cells were permeabilized with 0.5% Triton-X (T8787-50ML, Sigma-Aldrich) in 1xPBS for 10 minutes, 1% SDS (L6026-250G, Sigma-Aldrich) in 1xPBS for 15 minutes, and 0.1M HCl (SA56-1, Fisher chemical) for 5 minutes at room temperature. Three washes with 1xPBS were performed after each TritonX, SDS, or HCl treatment. Cells were then treated with 10 µg/mL RNaseA (EN0531, Thermo Scientific) in 1xPBS for one hour in a 37°C tissue culture incubator. After three 1xPBS washes, cells were treated with hybridization #1 buffer (0.1% (vol/vol) Tween-20 (P9416-50ML, Sigma-Aldrich), 50% (vol/vol) Formamide (S4117, MILLIPORE), 2xSSC (AM9763, Invitrogen)) for 35 minutes at room temperature. Meanwhile, a 22 mm x 22 mm coverslip (sc-363555, Santa Cruz Biotechnology) was coated with 150 µL of Sigmacote (SL2-100ML, Sigma-Aldrich) in a fume hood. After the hybridization #1 buffer was removed, 50 µL of hybridization #2 buffer (50% (vol/vol) Formamide, 2xSSC, 0.1% (vol/vol) Tween-20, 10% (vol/vol) Dextran Sulfate (BP1585-100, Fisher Scientific)) containing primary probes (∼500 ng) was added onto the fixed cells, and carefully covered with the Sigmacote-coated coverslip. The coverslip was heated at 90°C for 3 minutes, placed in an empty tip box filled with water, and incubated at 47°C in an incubator for 12 to 16 hours. Next day, the coverslip was washed with 2xSSC (pre-warmed at 47°C) in the 47°C incubator for 10 minutes. The 22 mm x 22 mm Sigmacote-coated coverslip was carefully removed, and the cells were washed with the pre-warmed 2xSSC in the 47°C incubator for 10 minutes. The cells were further washed with room temperature 2xSSC twice, followed by a post-fix buffer treatment (8% (vol/vol) formaldehyde, 2% (vol/vol) Glutaraldehyde (50-262-17, Fisher Scientific) in 1xPBS) for 1 hour at room temperature. The cells were then washed with 2xSSC three times. For Nanog and MYC gene primary probe hybridization, the temperature for overnight incubation and subsequent washes with pre-warmed 2xSSC was set to 42°C.

### ORCA: sequential hybridization, imaging, and image analysis

We performed sequential hybridization as previously described with some modifications ^23^. After primary probe hybridization, the coverslip was mounted on a FCS2 chamber (03060319-2-NH, Bioptechs). The FCS2 chamber was connected to a home-built fluidics system controlled by home-built software, and placed on the Leica DMi8 wide-field microscope imaging stage. First, 25% Ethylene Carbonate (EC; E26258-3KG, Sigma-Aldrich) in 2xSSC buffer containing 10 nM fiducial cy3 probes was perfused into the chamber and incubated for 15 minutes at room temperature. The sample was then washed with a wash buffer (30% (vol/vol) formamide in 2xSSC) and 2xSSC, and then an imaging buffer (0.25 mg/mL glucose oxidase (G2133-50KU, Sigma-Aldrich), 40 µg/ml catalase (02100402-CF, MP Biomedicals) and 9% (vol/vol) glucose (G8769-100ML, Sigma-Aldrich) in 2xSSC) was perfused in. Multi-position acquisition was manually set using the LAS X Leica microscope control software. The cy5 readout probes were prepared in a 96-well plate so that each well contains 100 nM adapter oligo, 110 nM cy5 readout oligo, and 300 nM displacement oligo to displace the previous round of hybridization, each in 750 µL of 25% EC buffer. The 96-well plate was covered with foil and placed on the fluidics together with ∼50 mL wash buffer, 2xSSC buffer and 20 mL imaging buffer. The imaging interval time was set using the home-built software. The automated fluidics was started first, and then imaging was started immediately after the first round of hybridization finished. Imaging was performed with a 63x/NA1.4 oil-immersion objective that generates 2048 pixel x 2048 pixel images at the scale of 103 nm/pixel, and using 200 nm z-steps to scan 6 µm vertically. For the fluorophore excitation, we used Lumencor LED illumination with 50% attenuation. The exposure times were 100 ms and 10 ms for cy5 and cy3, respectively.

To perform image analysis, we used ChrTracer3 developed by Alistair Boettiger lab^23^. The Leica-generated tif images were converted to dax format using a home-built software so that the files could be processed with ChrTracer3. The ChrTracer3 algorithm fits each fiducial and readout spot with a 3D gaussian to get 3-dimensional coordinates. The fiducial coordinates were used to correct 3-dimensional drift. Chromatin traces were obtained from the readout coordinates after drift correction. We also filtered readout spots by the median distance to other readout signals (cutoff 600 nm for AAVS1 and Nanog, 1000 nm for MYC) to eliminate non-specific background noise. To calculate the radius of gyration or perform t-SNE dimensionality reduction and k-mean clustering, traces were filtered to ensure segment detection efficiency above 50%, followed by linear interpolation to obtain complete 3D structures. The number of clusters for k-mean clustering (k=4) was empirically determined. To calculate the correlation between chromatin compaction and epigenetic memory or irreversible fate commitment, or perform statistical analysis such as Student’s *t*-test or Wilcoxon Rank Sum test, data from the same coverslip were used to minimize the impact of batch-to-batch variability when analyzing chromatin traces.

To compare chromatin traces at the MYC locus with Hi-C data, we used Hi-C data previously reported^22^. Normalized counts from the Hi-C data were exported as a matrix using JuiceBox^57^. The exported matrix was further analyzed using custom scripts in MATLAB.

### RNA FISH for reporter mRNA quantification

RNA FISH probes for the reporter mRNA were designed the same way as the ORCA primary probes, except they were designed based on the sense strand so that they hybridize to the mRNA. 3x10^4^ cells in 20 µL culture media were plated in the matrigel-coated PDMS multi-well device for 10 minutes and then 20 µL of 8% formaldehyde in 1xPBS was added to fix the cells. After removing the PDMS multi-well device, cells were permeabilized with 0.5% Triton-X in 1xPBS for 10 minutes at room temperature. After three 1xPBS washes, cells were treated with hybridization #1 buffer for 35 minutes at room temperature. After the hybridization #1 buffer was removed, 50 µL of hybridization #2 buffer containing 200 ng RNA FISH probes was added to the fixed cells, and carefully covered with a 22 mm x 22 mm Sigmacote-coated coverslip (as described in the ORCA protocol). The coverslip was placed in an empty tip box filled with water and incubated at 37°C in an incubator for 12 to 16 hours. Next day, the coverslip was washed with 2xSSC (pre-warmed at 37°C) in the 37°C incubator for 10 minutes. The 22 mm x 22 mm coverslip was carefully removed, and the cells were washed with the pre-warmed 2xSSC in the 37°C incubator for 10 minutes. The cells were further washed with room temperature 2xSSC twice. The RNA FISH signal was visualized using an adapter oligo that binds to the RNA FISH primary probes and has three tandem binding sites for the cy5 readout oligo. RNA FISH images were taken with a 63x/NA1.4 oil-immersion objective using 200 nm steps z-scanning. DAPI-staining images were taken after the RNA FISH imaging. Background was subtracted from the RNA FISH images using rolling ball background subtraction at the radius of 5 pixels. Then maximum intensity projection across the z-stack of images for a given XY position was used to generate single XY images. The number of RNA FISH spots was quantified using the spot detection algorithm in ImageJ with the prominence parameter set to 400. Cell masks were created using the StartDist plugin^58^ with the following parameters: ’modelChoice’:’Versatile (fluorescent nuclei)’, ’normalizeInput’:’true’, ’percentileBottom’:’20.0’, ’percentileTop’:’95.0’, ’probThresh’:’0.6’, ’nmsThresh’:’0.01’, ’outputType’:’Label Image’, ’nTiles’:’1’, ’excludeBoundary’:’2’, ’roiPosition’:’Automatic’. We filtered out cells with abnormal size, and then classified cells with less than 30 RNA FISH spots as silenced based on the FISH distribution for cells with KRAB recruited for 5 days (+dox) that are fully silenced by flow cytometry.

### CUT&RUN for detection of histone modifications

CUT&RUN to profile histone modifications was performed as previously described, using CUTANA ChIC/CUT&RUN Kit (14-1048, EpiCypher) and H3K9me3 antibody (C15410193, Diagenode)^21^. An input of 5x10^5^ HEK293T cells was processed according to the manufacturer’s protocol. The double integrant clone (B9) was used for this experiment. Digitonin was used at a final concentration of 0.025% for nuclear permeabilization. Dual-indexed sequencing libraries were made using the NEBNext Ultra II DNA Library Prep Kit for Illumina (E7645S, New England Biolabs) or CUTANA™ CUT&RUN Library Prep Kit with Primer Set 1 (14-1001, EpiCypher), and library concentrations were quantified with the Qubit 1 X dsDNA HS Assay Kit (Q33231, Invitrogen). Library fragment sizes were checked with Agilent 2100 Bioanalyzer through the Protein and Nucleic Acid Facility at Stanford University School of Medicine. For sequencing, a NextSeq System (Illumina) was used. A custom human genome (version hg38) with the reporter integration was constructed using bowtie2-build. Paired-end alignment was performed with a bowtie2 command, and duplicate reads were removed using Picard. Bedgraph files were generated from the reads, normalized by dividing by total counts in each sample, and reported as counts per million. Further analyses and visualization were performed using MATLAB. For H3K9ac and H3K27ac quantification, reads aligned to human EF1alpha promoter were excluded because they also include signal from the endogenous EF1alpha promoter or the EF1alpha promoter that drives expression of rTetR-KRAB in addition to the target EF1alpha in the reporter (pEF).

### EM-seq for detection of DNA methylation

Enzymatic methyl-seq (EM-seq) was performed to profile DNA methylation. Briefly, 1x10^6^ HEK293T cells were collected and their gDNA was extracted using the NEB Monarch kit (T3010S, NEB) according to the manufacturer’s protocol. gDNA was subjected to restriction enzyme digestion with XbaI and AccI for 1 hour at 37°C. Following SPRIselect purification (B23318, Beckman-Coulter), digested DNA was converted using New England Biolabs’ Enzymatic Methyl-seq Kit (E7120S, NEB) to detect selectively oxidized unmethylated cytosines. After conversion, 3 PCRs were performed to selectively amplify a segment of 771 bp of reporter DNA overlapping the pEF promoter. First, NEB Q5U polymerase (M0515, NEB) was used according to manufacturer’s specifications to amplify reporter DNA from the pool of gDNA while maintaining the identity of converted uracils. Subsequently, amplified DNA was amplified again using NEB Q5 Ultra II to add Illumina Read1 and Read2 overhangs (M0544, NEB). Finally, unique sequencing indices were added to each sample in a third PCR reaction also using NEB Q5 Ultra II. For sequencing, a MiSeq System (Illumina) was used with a 600v3 kit, reading 374 cycles in read1 and 235 cycles in read2. BWAmeth was used for methylation-specific alignment to the reporter, and samtools was used to convert raw alignment files to indexed .bam. Bulk CpG methylation levels were calculated per site using MethylDackel (https://github.com/dpryan79/MethylDackel). Custom analyses were used (dSMF-footprints_optional_clustering.py) to produce single-molecule methylation heatmaps of aligned reporter molecules. Further analyses and visualization were performed using MATLAB.

### Mouse ES cell culture and differentiation

Mouse embryonic stem cells CASTx129 were cultured in 2i+LIF serum-free media (stem cell media). The media was made by mixing 0.5X DMEM/F12 (11320-033, Gibco), 0.5X Neurobasal Medium (21103049, Thermo Scientific), 1X GlutaMAX Supplement (35050061, Thermo Fisher Scientific), 0.05% Bovine Albumin Fraction V (7.5% solution) (15260037, Thermo Scientific), 100uM 2-Mercaptoethanol (50 mM) (31350010, Thermo Scientific), 0.5X N-2 Supplement (100X) (17502-048, Thermo Scientific), 1X PEN/STREP 100X (15-140-122, Thermo Fisher Scientific), and 0.5X B-27 Supplement (50X) serum free (17504044, Thermo Scientific), supplemented with 1X 0.1mg/mL LIF 10000X (130-095-777, Miltenyi Biotec), 1X 5mM PD0325901 5000X (1408, Axon Med Chem) and 1X 15mM CHIR99021 5000X (1386, Axon Med Chem). A culture dish was pre-coated with 0.0015% poly-L-Ornithine (P4957-50ML, Sigma-Aldrich) for one hour in a 37°C tissue culture incubator followed by 10 µg/mL Laminin (L2020-1MG, Sigma-Aldrich) coating in the same way. For passaging, cells were trypsinized for 5 minutes with 0.25% Trypsin-EDTA (252-000-56, Fisher Scientific), spun down after deactivation with equal amount of the stem cell media, and resuspended in fresh media because the leftover of trypsin inhibits cell adhesion. Cells were plated at a density of 3x10^5^ cells/well in a 6-well plate. The stem cell medium was exchanged every day and cells were passaged onto a new plate every other day.

To differentiate mouse ES cells, stem cell media without LIF, PD0325901 and CHIR99021 were used as the differentiation media (N2B27 media). Cells were plated at a density of 1x10^5^ cells/well in 6-well plates in 2mL differentiation media. After 1, 2 or 3 days of differentiation, 1x10^5^ trypsinized cells were spun down and frozen at -80°C for qPCR. For subsequent ORCA measurements, 2x10^5^ trypsinized cells were spun down and resuspended in 100 µL 4% formaldehyde in 1xPBS for 10 minutes. These fixed cells were then spun down and resuspended in 30 µL of 70% ethanol and stored at -20°C until they were used for ORCA measurements. For use in the colony-based irreversibility assay, 2x10^3^ trypsinized cells were spun down and resuspended in 100 µL culture media, spun down again and resuspended in 200 µL stem cell media. Of these, 500 cells in 50 µL were added to 500 µL stem cell media in 24-well plates coated with 0.0015% poly-L-Ornithine and 10 µg/mL Laminin and grown further for 5 days.

### Mouse ES cell differentiation irreversibility assay based on colony formation

Mouse embryonic stem cells replated in the stem cell culture media after differentiation (see Mouse ES cells culture and differentiation section above) were fixed by adding 500 µL 8% formaldehyde in 1xPBS followed by 1xPBS wash twice. Cells were incubated with 1 µg/mL DAPI (AC202710500, Fisher Scientific) for 10 minutes, and each well was scanned with a 4x objective on the Leica DMi8 . To estimate the number of colonies that originated from each individual dedifferentiated cell, images were processed as follows using ImageJ: a median filter with a radius of 4 was applied to all images, then rolling ball background subtraction at a scale of 20 was performed. A Gaussian filter with sigma of 4 was applied, then the images were binarized by Li thresholding. Pixel clusters were filtered to keep only the ones with a size of 40-10000 pixels. Watershed was applied to further separate colonies from background noise (especially those close to the edge of the well). A second pixel cluster filtering was performed to select clusters with sizes between 40-2500 pixels and with circularity of 0.7 to 1.0, which were defined as colonies.

### RT-qPCR of fate marker genes

For each differentiation condition, 1x10^5^ cells were spun down and stored at -80°C as a cell pellet after removing culture media. After thawing, total RNA was extracted using the RNeasy Mini Kit (74106, Qiagen). After quantifying single-strand RNA concentration using Nanodrop, cDNA synthesis was performed using iScript Reverse Transcription Supermix (1708840, Bio-Rad). For qPCR, SsoAdvanced Universal SYBR Green Supermix (1725270, Bio-Rad) was used. We used previously reported primer sets^32^. CFX Connect (Bio-Rad) was used for the qPCR readout. The Tbp gene was used as an internal normalization control, and mRNA levels for each gene relative to Tbp were defined as 2^(Cq_Tbp-Cq_gene)^, then normalized to the day0 mRNA level so that the average mRNA at day0 is equal to 1.

### ORCA: mouse ES cells

Before palting the fixed ES cells suspensions onto a 40 mm coverslip, 5 µL 0.01% poly-L-Lysine was added in the PDMS multi-well device (each with a 3 mm inner diameter) pressed onto a 40 mm coverslip and completely air-dried for one hour in a tissue culture hood. 7.5 µL of ES cells suspensions fixed in 70% ethanol (as described in the see Mouse ES cell culture and differentiation section) was added into the PDMS wells and incubated for 10 minutes at room temperature. The 70% ethanol was removed and the PDMS multi-well device was carefully peeled off. After air-drying for 1 minute, cells were fixed with 4% formaldehyde in 1xPBS for 10 minutes. The rest of the primary probe hybridization protocol was the same as described in the ORCA protocol: primary probe hybridization section. We performed sequential hybridization as described above. To make this chromatin tracing comparable to the experiment performed at the synthetic reporter gene, we targeted a 60 kb region around the Nanog gene promoter at a resolution of 5 kb, and three additional 5kb-long segments upstream and downstream at 15 kb intervals.

### 3D random-walk polymer model fitting

We used a Bayesian optimization algorithm to fit a locally-compacted 3D random-walk polymer model to the experimental data (Extended Data Fig. 9). Analysis was performed using python on Google Colaboratory Pro. The Cupy package^59^ was used to parallelize the computation on multiple GPUs, and the Optuna package^60^ was used for Bayesian optimization. In each trial, 50000 3D random-walk polymers were generated with randomly sampled step sizes. The parameter (step-size) search range was set to 70 nm to 210 nm. The TPE sampler was used to optimally sample the random-walk step size, with the prior_weight parameter set to 1000. The objective function to be minimized was defined as the sum of squared errors between the median distance maps measured experimentally and the median distance maps computed from the simulated 3D random-walk polymers. Trials were repeated 300 times, and the parameter set (of step-sizes at each position) with the least error was chosen as the best fit. 10 independent optimizations were performed and the median of the best fit parameter sets was used for further calculations (eg. radius of gyration).

### Statistics and Reproducibility

Each experimental condition has at least two replicates. Each replicate includes 456 to 1227 traces (Supplementary Table 1). The overall correlation coefficient between replicates is 0.76 (Extended Data Fig. 3). All statistical analyses were performed using the data on the same coverslip (i.e. same experiment ID in Supplementary Table 1).

## Data Availability

CUT&RUN data has been deposited on the Gene Expression Omnibus (GEO) database, and it is currently under review. All processed ORCA data will be publicly available through the 4DN portal.

## Code Availability

Custom codes associated with this study have been uploaded to Github (https://github.com/bintulab/ORCA_Fujimori_2023).

## Author contributions

T.F. and L.B. conceptualized the project. T.F. designed and performed all experiments and analyses except EM-seq. C.R. helped with the VP64 experiment and made the DNMT3B cell line. C.R. and A.R.T. performed the EM-seq experiment. M.M.H. and A.R.T. designed EM-seq primers. B.R.D. built the EM-seq analysis pipeline. W.J.G. supervised the EM-seq workflow development. A.H. validated the ORCA probe at the MYC locus. D.L. synthesized and validated the ORCA probe at the Nanog locus together with A.H.. J.S. made the HDAC4 plasmid. A.N.B. designed the ORCA probe at the MYC locus and the Nanog locus, and helped build ORCA on a commercial wide-field microscope. L.B. supervised the project. T.F. and L.B. wrote the manuscript with input from all authors.

## Acknowledgements

We thank all current and past members of the Bintu and Boettiger labs for the fruitful discussions and suggestions. We thank Raeline Valbuena for their input on designing the droplet digital PCR experiment. We thank Sedona Murphy for numerous suggestions on improving DNA FISH. We thank Jude Lee and Liang-Fu Chen for advice on mouse ES cell culture. We thank Minhee Park for technical assistance with performing the MYC ORCA controls on the Boettiger lab custom microscope. We thank Bogdan Bintu for suggestions on building the fluidics system. We thank the Michael Bassik lab for access to the NextSeq. This work was supported by the NIH 4D Nucleome U01DK127419 (L.B., A.N.B.), JSPS Overseas Research Fellowships (T.F.), NSF Graduate Research Fellowship DGE-1656518 (B.R.D.).

