## Supplemental materials for "Single-cell chromatin state transitions during epigenetic memory formation"

### Extended Data Figures

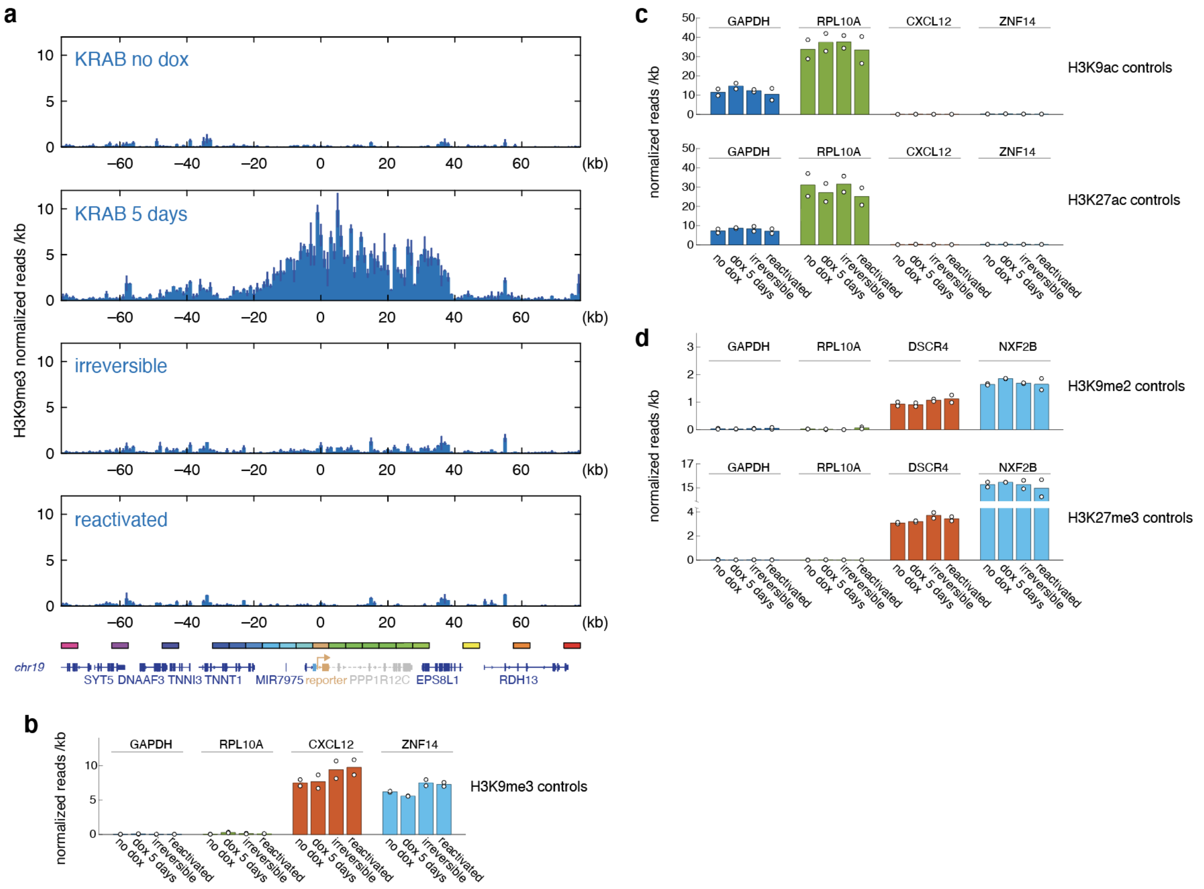


**Extended Data Fig. 1 | Changes in epigenetic modifications upon KRAB recruitment.** (a) Genome traces showing normalized number of reads after CUT&RUN against H3K9me3 as a function of distance around the reporter integration site (at 0 kb), in no dox control cells, upon KRAB recruitment for 5 days, in irreversibly silenced cells, or reactivated cells. Irreversibly silenced cells and reactivated cells were obtained by treating with dox for 5 days, washing out dox for 22 days, sorting the population with mCitrine reporter OFF or ON respectively, and growing the cells for the CUT&RUN assay. Bar plots show the average from two replicates, blue lines show maximum and minimum values. (b) Quantification of the normalized H3K9me3 CUT&RUN reads at negative (GAPDH, RPL10A) and positive (CXCL12, ZNF14) control gene loci. (c) Quantification of the normalized H3K9ac or H3K27ac CUT&RUN reads at positive (GAPDH, RPL10A) and negative (CXCL12, ZNF14) control gene loci. (d) Quantification of the normalized H3K9me2 or H3K27me3 CUT&RUN reads at negative (GAPDH, RPL10A) and positive (DSCR4, NXF2B) control gene loci. Bars show the average from two replicates, dots represent individual replicates. All CUT&RUN experiments were done with the B9 clone (see Extended Data Fig. 4). Note that epigenetic modification enrichment at positive and negative control genes are consistent across conditions.

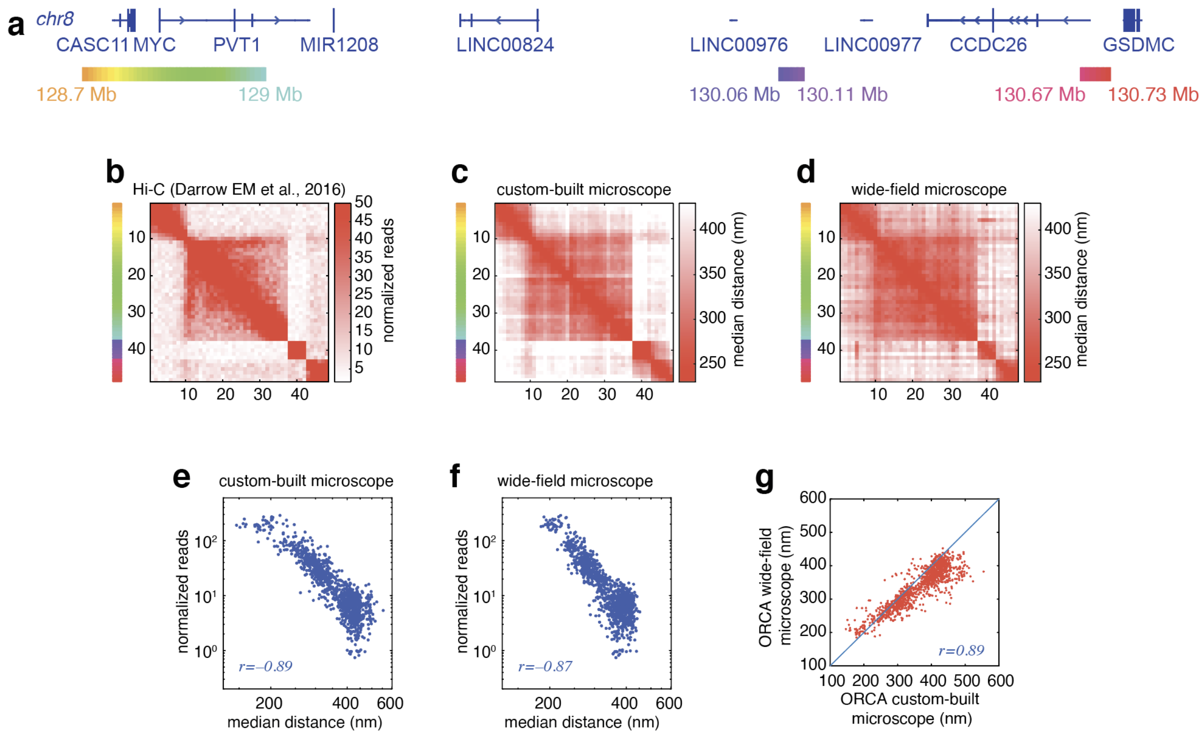


**Extended Data Fig. 2 | Validation of the ORCA system built on a wide-field microscope.**

(a) ORCA probe design at the MYC locus. (top) Gene annotations. (bottom) Locations of ORCA probes are shown in different colors. Each probe set targets a 10 kb segment. (b) A heatmap showing the pairwise contact frequency as computed from bulk Hi-C normalized reads at the MYC locus (data from Darrow EM and Huntley MH *et al.*, *PNAS*, 2016[22]). (c) Median distance map between pairs of segments across the MYC locus measured using ORCA on a custom-built microscope that has been used for previous publications (Boettiger lab[18]). (d) Median distance map of ORCA on the wide-field microscope used in this study (Bintu lab). (e) Correlation between Hi-C normalized reads and ORCA median distances for the MYC locus measured on the custom-built microscope. (f) Correlation between Hi-C normalized reads and ORCA median distances measured on the wide-field microscope. Linear correlation between Hi-C number of reads and median distances in a log scale was observed as reported before[17]. (g) Correlation between median distances measured by ORCA on a custom-built microscope vs. the wide-field microscope used in this study.

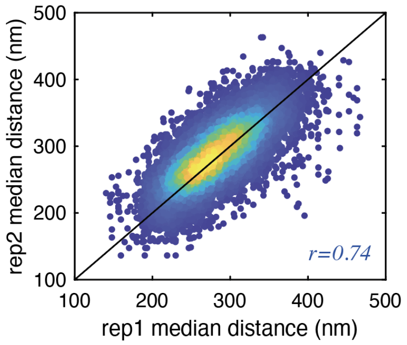


**Extended Data Fig. 3 | Reproducibility of the median distances measured by ORCA at the AAVS1 locus.** Median distances of all pairwise segments from all pairs of replicates across all datasets from all HEK293T cell lines derivatives of the B9 clone reported in this study are superimposed. In the case of multiple replicates, all combinations of replicate pairs are all plotted (eg. rep1/rep2, rep2/rep3, rep1/rep3, etc.). The overall correlation coefficient is 0.76.

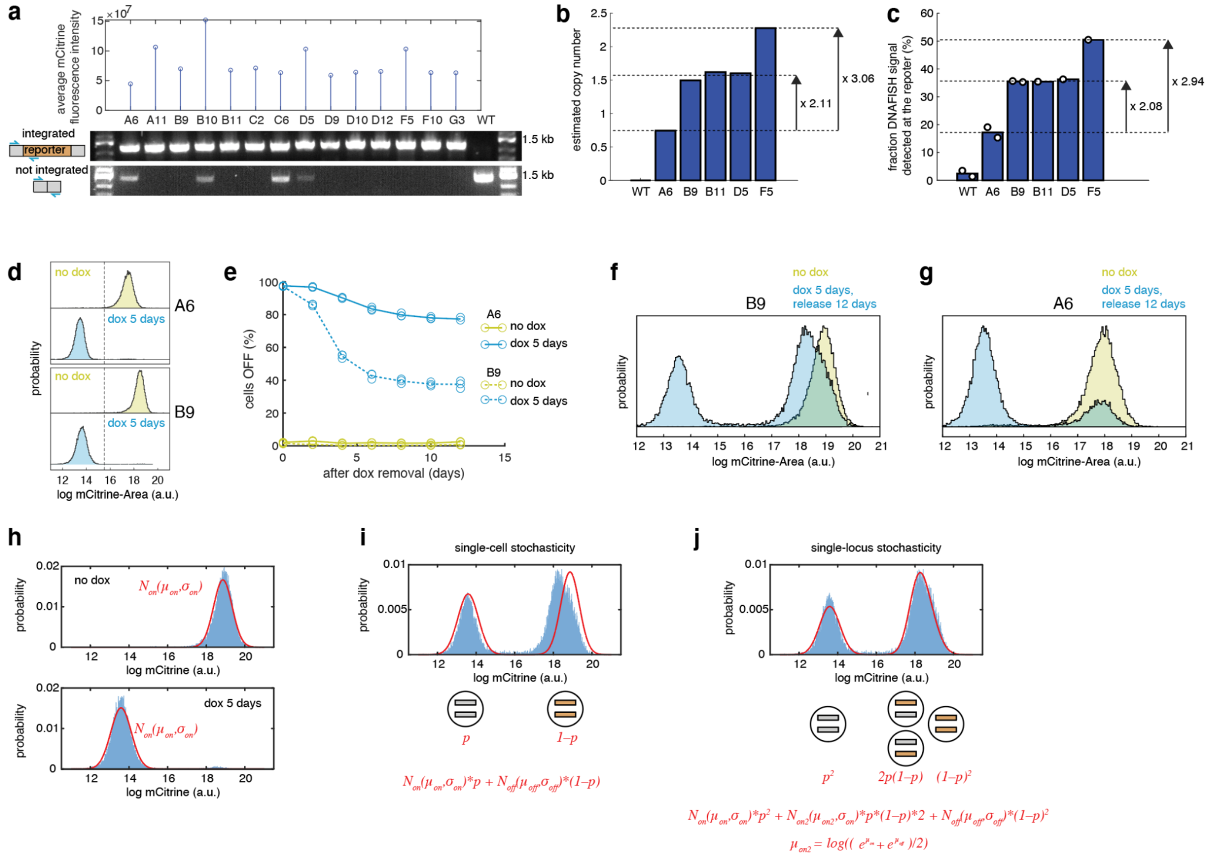

**Extended Data Fig. 4 | Epigenetic memory for cell clones with different numbers of reporter integrations.**

(a) We used limiting dilution to select single-cell reporter clones (A6-G3), expanded them, and characterized them to determine the number of reporter integrations at the AAVS1 locus on chromosome 19 which exists in 3 copies in HEK293T cells. (top) Average mCitrine fluorescence intensity for the single clones as measured by flow cytometry. (bottom) Genomic PCR results with primers that detected integrated reporter or the WT sequences (not integrated). (b) Estimated copy number of the reporter gene relative to the Albumin gene measured by droplet digital PCR. (c) The percentage of chromosomes with positive DNA FISH readout signal at the reporter out of all chromosomes 19 detected by the fiducial signal. (d) Flow cytometry histograms of the mCitrine fluorescence intensity for the single reporter integrant clone (A6, top) or the double reporter integrant clone (B9, bottom) after integration of the rTetR-KRAB construct by lentivirus and treatment with no dox (yellow) or dox 5 days (blue). (e) Time-course showing the percentage of cells with mCitrine OFF for the A6 or B9 cell lines during the KRAB release period following 5 days of KRAB recruitment. (f) Histograms of the reporter mCitrine fluorescence intensity of the B9 line after 5 days with dox and 12 days of dox removal (blue) versus no dox (yellow) as measured by flow cytometry. Note that reactivated cells show lower mCitrine expression levels compared to no dox, which is not observed in the single integrant A6 clone in g. (g) Histograms of the reporter mCitrine fluorescence intensity of the A6 line after 5 days with dox and 12 days of dox removal (blue) versus no dox (yellow) as measured by flow cytometry. (h) Histograms of mCitrine fluorescence for the double reporter B9-KRAB cell line were fitted with a normal distribution (imposing a ± 3 sigma cutoff) for (top) cells with no dox or (bottom) treated with dox for 5 days. (i) Fluorescence intensity distribution of the mCitrine reporter after reactivation in the double reporter B9-KRAB cell line (blue) was fitted with a probability distribution (red) that assumes stochasticity happens at the cell level (i.e. if a given reporter activates in a cell, the other reporter also reactivates in that cell, as shown in the schematic on the bottom). This cell stochasticity predicts a higher fluorescence distribution (red) than measured (blue). (j) Fluorescence intensity distribution of the mCitrine reporter after reactivation in the double reporter B9-KRAB cell line (blue) was fitted with probability distribution assuming stochasticity happens at the individual locus level (i.e. in a given cell, each integrated reporter can activate independently of the reporter integrated on another chromosome, as shown in the schematic on the bottom). Note that the independent locus stochasticity model in j shows better fitting than the cell-driven stochasticity model in i.

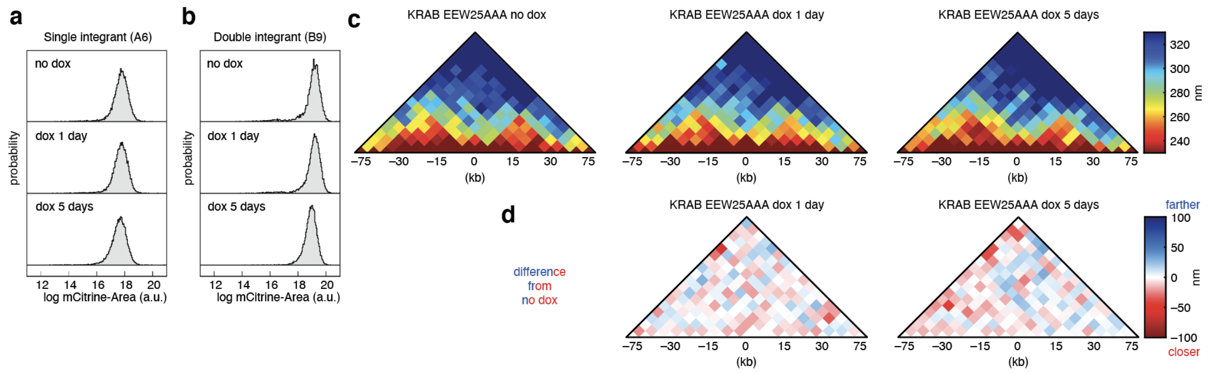


**Extended Data Fig. 5 | Chromatin compaction is not observed after KRAB EEW25AAA mutant recruitment.** (a) Flow cytometry histograms of the reporter mCitrine fluorescence intensity for the single integrant clone (A6) with no dox, dox 1 day, or dox 5 days. (b) Flow cytometry histograms of the reporter mCitrine fluorescence intensity of the double integrant clone (B9) with no dox, dox 1 day, or dox 5 days. (c) Median distance maps from chromatin traces upon KRAB EEW25AAA mutant recruitment for 1 day or 5 days and no dox control. (d) Subtracted median distance maps upon 1 day or 5 days KRAB EEW25AAA recruitment compared to the no dox control. Two replicates are averaged.

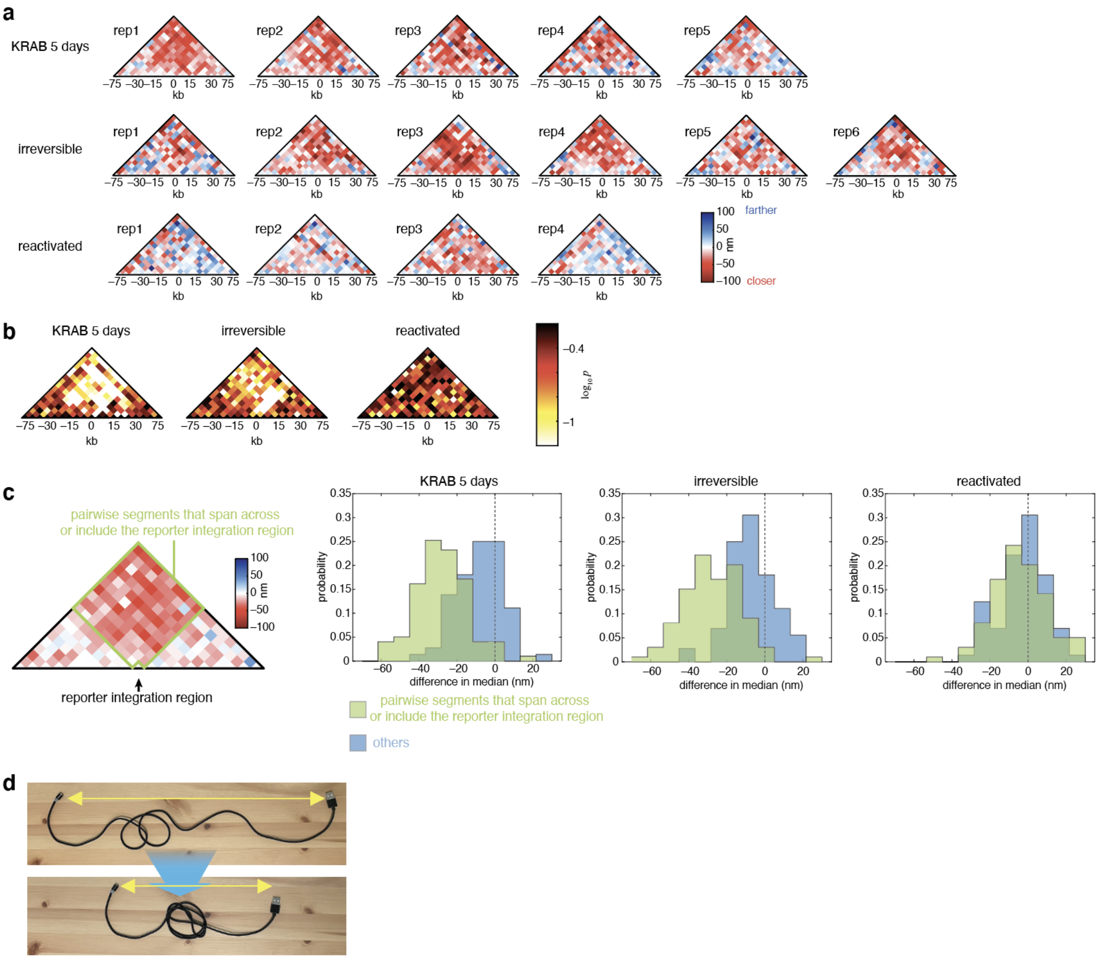


**Extended Data Fig. 6 | Reproducibility of chromatin compaction, and localized chromatin compaction.** (a) Subtracted median distance maps compared to no dox controls for each replicate that contributes to the average distance maps in Fig. 1e,f. (b) Statistical analysis of pairwise distances showing log-transformed *p* values averaged across replicates calculated using the Wilcoxon Rank Sum test for the indicated samples compared to the no dox control. (c) (left) a green box highlights pairwise segments that span across the reporter integration region (contain the reporter site between them). (right) Histograms of subtracted median distances between segments that span across or include the reporter integration region (green box in the left figure), and other pairwise segments (blue). (d) Visual interpretation of locally compacted chromatin. “Compaction” at the center of a cord makes end-to-end distance shorter (yellow arrows).

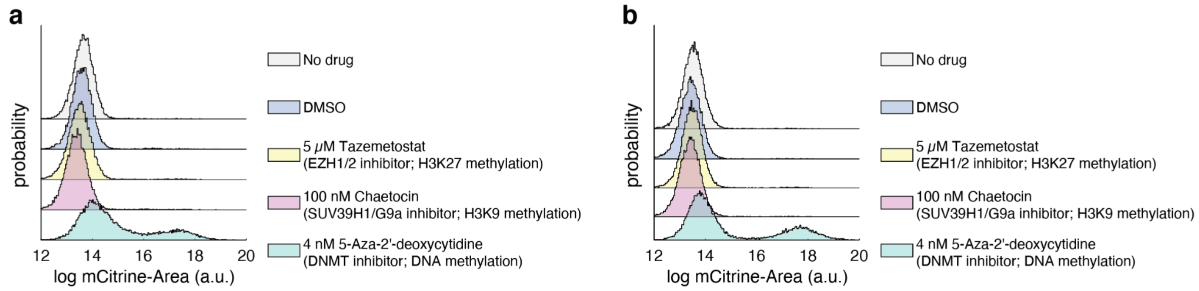


**Extended Data Fig. 7 | A DNA methyltransferase inhibitor reactivates irreversibly silenced cells.** (a) Flow cytometry histograms of mCitrine intensity for irreversibly silenced cells (generated as described in Fig. 1d) upon treatment with the H3K27 methyltransferase inhibitor Tazemetostat, H3K9 methyltransferase inhibitor Chaetocin, or DNA methyltransferase inhibitor 5-Aza-2'-deoxycytidine for 4 days. (b) Flow cytometry histograms of mCitrine intensity for irreversibly silenced cells 8 days after inhibitors removal. Experiments were performed with the B9 clone.

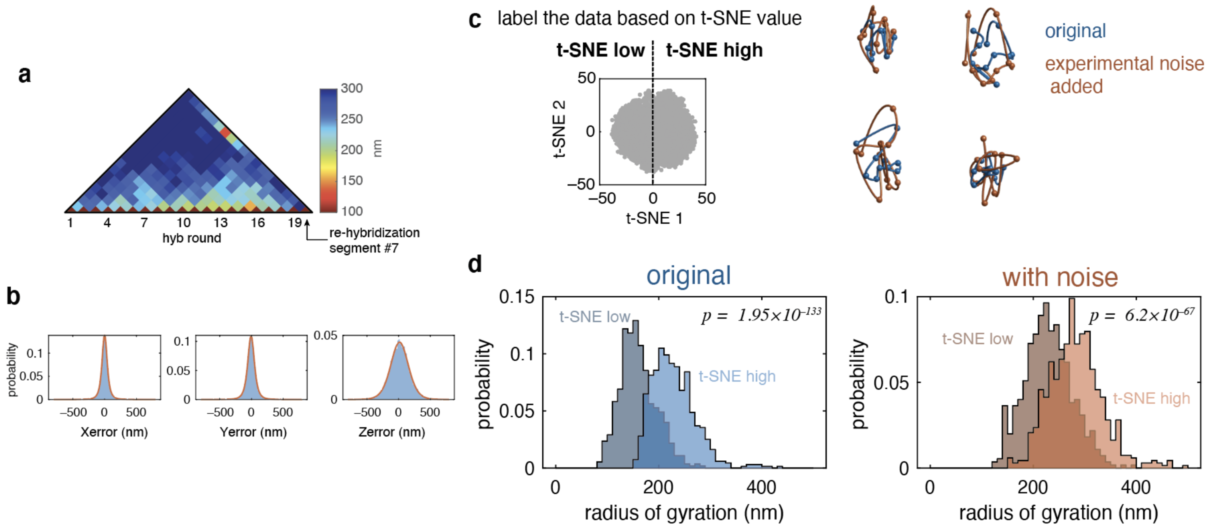


**Extended Data Fig. 8 | The effect of experimental noise on heterogeneity in the t-SNE dimensionality reduction analysis.** (a) A median distance map for B9-KRAB no dox cells with segment #7 imaged during round 7 of imaging and re-hybridized at the end of chromatin tracing (black arrow). Note that the distance between segment #7 and the re-hybridized segment #7 is the smallest across all pairwise distances (red). (b) The difference between 1$^{st}$ and 2$^{nd}$ hybridization of segment #7 in each direction in 3D (blue). The distributions were fitted with a t-location scale distribution (red). (c) (left) Samples were classified into t-SNE high (t-SNE 1 > 0) or t-SNE low (t-SNE 1 < 0). (right) Simulated experimental noise was sampled from the fitted distribution in b, then added to each chromatin trace (original traces are in blue and traces with simulated experimental noise are in red). (d) The distribution of radius of gyration for the t-SNE low group (dark color) and the t-SNE high group (light color) before (blue) or after (red) adding experimental noise. The p-values were calculated using the Wilcoxon Rank Sum test.

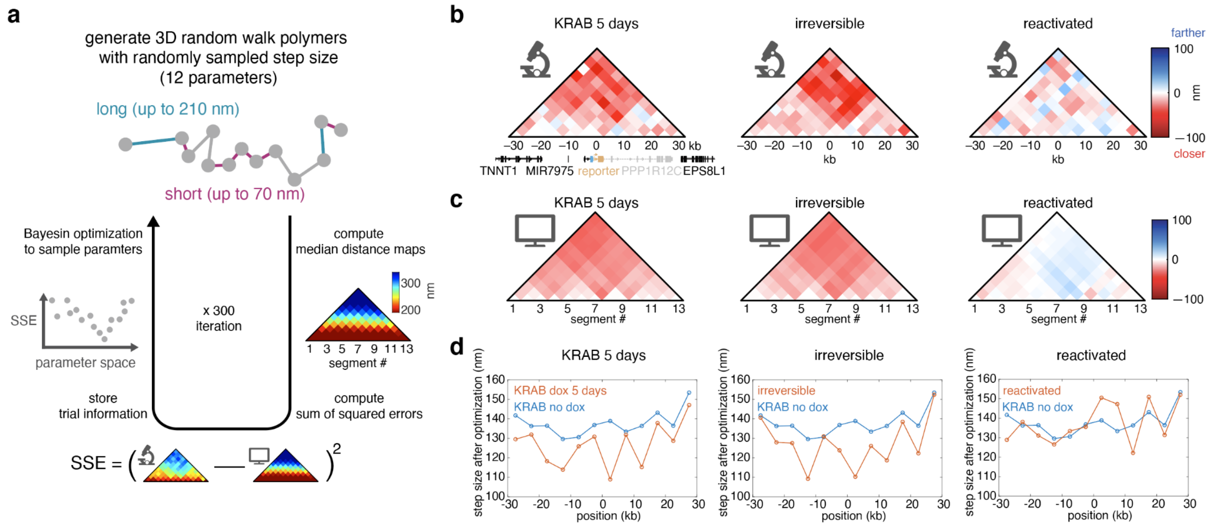


**Extended Data Fig. 9 | Bayesian optimization to fit a locally-compacted 3D random-walk polymer model to experimental data.** (a) A schematic of the pipeline for Bayesian optimization to fit a 3D random-walk polymer model to experimental data (Methods). (b) Subtracted median distance maps from the experimental data used for fitting. (c) Subtracted median distance maps generated using the fitted random-walk polymer model. (d) Fitted step sizes at each genomic position.

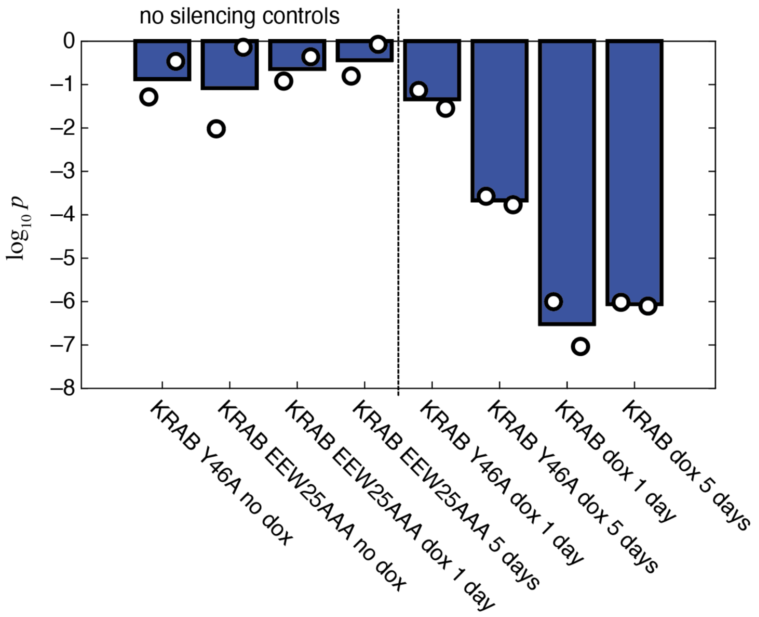


**Extended Data Fig. 10 | Statistical analysis of the medians radii of gyration.** Wilcoxon Rank Sum test was carried out as a measure of the difference in the median radius of gyration for each condition compared to WT KRAB no dox cells. Bar plots show the average. Dots represent two biological replicates.

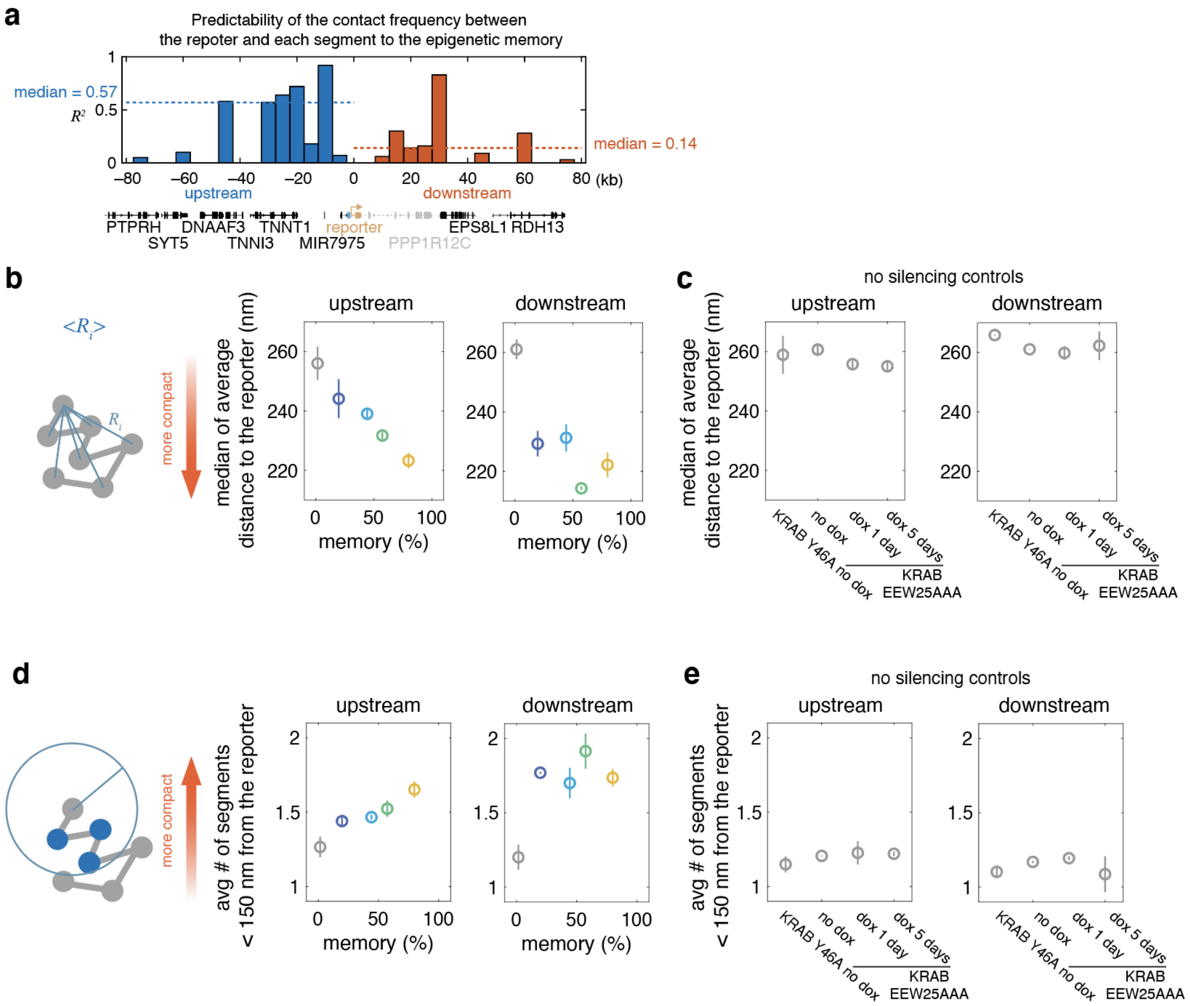


**Extended Data Fig. 11 | The upstream region is more predictive of epigenetic memory than the downstream region for the AAVS1 reporter system.** (a) Predictability of the contact frequency between the reporter and each segment defined as the $R^2$ value after linear regression as in Fig. 4f. Dashed lines show the median of the $R^2$ values across segments in the upstream region or the downstream region. (b) Median of the average distance from the reporter towards (left) upstream or (right) downstream plotted against different memory levels (%). (c) No silencing controls for b. (d) Average number of segments in (left) upstream or (right) downstream region within 150 nm from the reporter plotted against different memory levels (%). (e) No silencing controls for d.

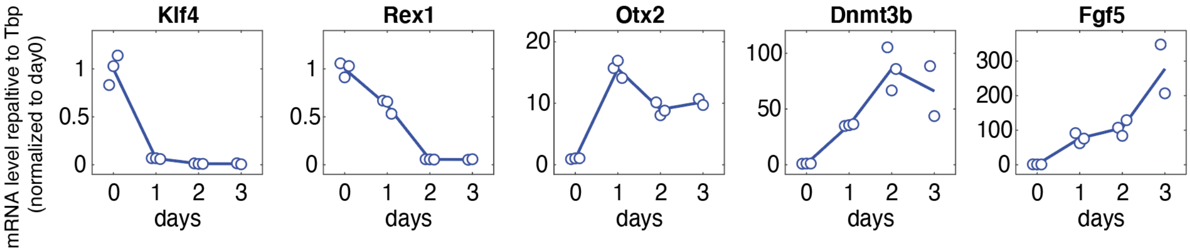


**Extended Data Fig. 12 | Gene expression changes for cell markers upon mouse ES cell differentiation.** Marker genes for the pluripotent state (Klf4, Rex1) and the differentiated state (Otx2, Dnmt3b, Fgf5) were quantified by RT-qPCR. The mRNA levels were quantified relative to Tbp, then normalized to the average mRNA level at day 0. Line plots show the average across two or three biological replicates.

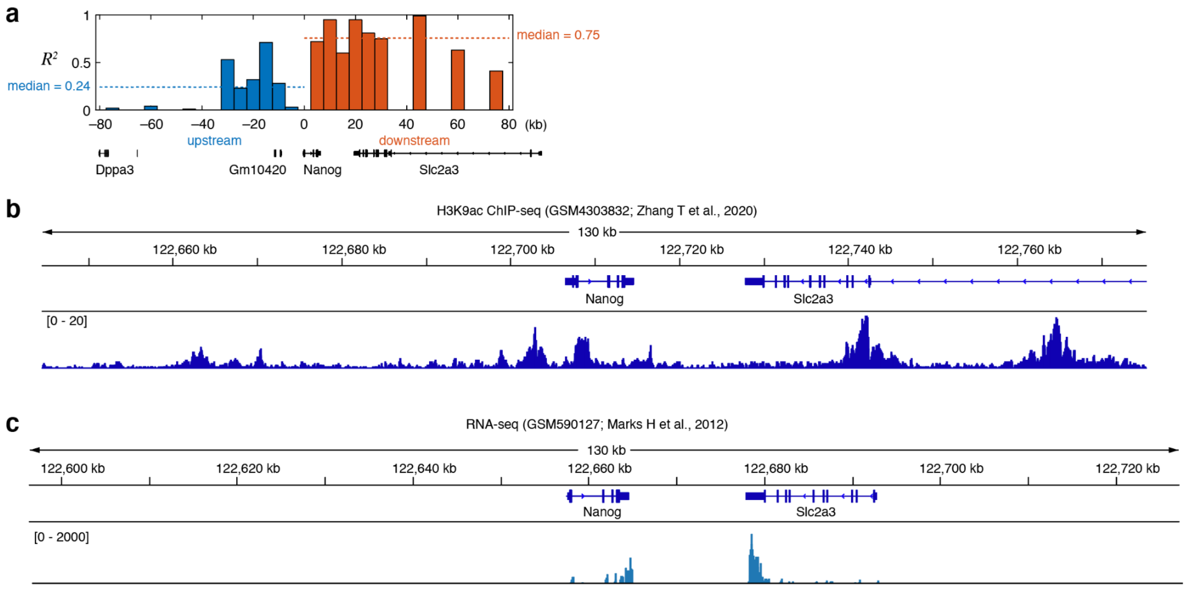


**Extended Data Fig. 13 | Contact frequency between the Nanog locus and the downstream region is more predictive of fate commitment in mouse ES cell differentiation.** (a) Predictability of the contact frequency between the Nanog locus and each segment defined as the $R^2$ value after linear regression, as in Fig. 5g. Dashed lines show the median of the $R^2$ values across segments in the upstream region or the downstream region. (b) Genome traces showing the normalized number of reads after ChIP-seq against H3K9ac in mouse ES cells. (Data from Zhang T *et al.*, *Genome Biol.*, 2020[53], GEO accession number: GSM4303832. Sequences were aligned to the mouse mm10 reference genome.) (c) RNA-seq in mouse ES cells (Data from Marks H *et al.*, *Cell*, 2012[54], GEO accession number: GSM590127. Sequences were aligned to the mouse mm9 reference genome.

| Experimental condition | replicate | experiment ID | Number of traces |
|---|---|---|---|
| HDAC4 dox 1 day | 1 | 3 | 840 |
| HDAC4 dox 1 day | 2 | 4 | 890 |
| HDAC4 dox 1 day | 3 | 6 | 922 |
| HDAC4 no dox | 1 | 3 | 835 |
| HDAC4 no dox | 2 | 4 | 944 |
| HDAC4 no dox | 3 | 6 | 639 |
| KRAB dox 1 day | 1 | 1 | 883 |
| KRAB dox 1 day | 2 | 2 | 967 |
| KRAB dox 1 day | 3 | 4 | 566 |
| KRAB dox 1 day | 4 | 5 | 456 |
| KRAB dox 1 day | 5 | 6 | 730 |
| KRAB dox 5 days | 1 | 1 | 788 |
| KRAB dox 5 days | 2 | 2 | 890 |
| KRAB dox 5 days | 3 | 3 | 604 |
| KRAB dox 5 days | 4 | 4 | 495 |
| KRAB dox 5 days | 5 | 6 | 688 |
| KRAB no dox | 1 | 1 | 989 |
| KRAB no dox | 2 | 2 | 1126 |
| KRAB no dox | 3 | 3 | 777 |
| KRAB no dox | 4 | 4 | 1081 |
| KRAB no dox | 5 | 5 | 482 |
| KRAB no dox | 6 | 6 | 552 |
| KRAB no dox | 7 | 7 | 704 |
| KRAB no dox | 8 | 9 | 1025 |
| KRAB Y46A dox 1 day | 1 | 1 | 954 |
| KRAB Y46A dox 1 day | 2 | 2 | 936 |
| KRAB Y46A dox 5 days | 1 | 1 | 1020 |
| KRAB Y46A dox 5 days | 2 | 2 | 741 |
| KRAB Y46A no dox | 1 | 1 | 1078 |
| KRAB Y46A no dox | 2 | 2 | 1108 |
| KRAB EEW25AAA dox 1 day | 1 | 1 | 1124 |
| KRAB EEW25AAA dox 1 day | 2 | 2 | 1123 |
| KRAB EEW25AAA dox 5 days | 1 | 1 | 1088 |
| KRAB EEW25AAA dox 5 days | 2 | 2 | 1089 |
| KRAB EEW25AAA no dox | 1 | 1 | 1195 |
| KRAB EEW25AAA no dox | 2 | 2 | 1227 |
| VP64 dox 1 day | 1 | 7 | 480 |
| VP64 dox 1 day | 2 | 8 | 773 |
| VP64 no dox | 1 | 7 | 546 |
| VP64 no dox | 2 | 8 | 641 |
| KRAB irreversible | 1 | 3 | 751 |
| KRAB irreversible | 2 | 4 | 621 |
| KRAB irreversible | 3 | 5 | 504 |
| KRAB irreversible | 4 | 6 | 816 |
| KRAB irreversible | 5 | 7 | 515 |
| KRAB irreversible | 6 | 9 | 731 |
| KRAB reactivated | 1 | 3 | 888 |
| KRAB reactivated | 2 | 4 | 921 |
| KRAB reactivated | 3 | 5 | 668 |
| KRAB reactivated | 4 | 6 | 833 |

**Supplementary Table 1**

Summary table of the number of replicates and traces for each experimental condition.
